# β_3_-adrenergic browning of pericardial adipose tissue controls cardiac function

**DOI:** 10.1101/2025.09.30.679591

**Authors:** Henver S. Brunetta, Bridget Coyle-Asbil, Alexa N. King, Anna E. Kupraty, Geneviève J. DesOrmeaux, Rachel M. Handy, Pierre-Andre Barbeau, Aleah J. Kirsch, Thomas Pulinilkunnil, Jean-Francois Legare, Petra Kienesberger, Jeremy A. Simpson, Keith R. Brunt, Graham P. Holloway

## Abstract

Dysfunctional adipose tissue (AT) is strongly linked to the development of cardiovascular diseases (CVD). Accumulation of AT around vital organs is detrimental to their respective function and overall health. Although there is strong evidence linking the accumulation of pericardial AT with CVD development, a comprehensive investigation on the adaptation of pAT in obesity is scarce. Here, by applying pair-wise bottom-up proteomics in pAT of humans and mice, we found pAT presents a browning signature, as demonstrated by enrichment of mitochondria, presence of UCP1, and greater metabolic capacity compared to subcutaneous AT. In mice fed a high-fat diet or obese patients, the pAT undergoes whitening, characterized by adipocyte hypertrophy, reduced mitochondrial content, respiratory capacity, and UCP1 levels. Lipectomy of pAT from obese mice decreased pathological ventricular hypertrophy. Conversely, selective β_3_-adrenergic agonist treatment rescued pAT browning status and is associated with improved heart structure and function, including ventricular thickness, and fibrosis in obese mice. Importantly, lipectomy of pAT abrogated the positive effects of β_3_-adrenergic agonism in cardiac function of obese mice. Altogether, our work positions pAT as a mechanistic driver of obesity-related cardiac dysfunction and establish β_3_-adrenergic-mediated browning of pAT as a novel therapeutic treatment strategy.

## Introduction

Obesity is one of the strongest independent risk factors for the development of cardiovascular disease (CVD), the leading cause of death worldwide^1^. Obesity is characterized by adipose tissue (AT) dysfunction, marked by adipocyte hypertrophy, impaired endocrine function, disrupted substrate management, and mitochondrial dysfunction^2,3^. AT in direct contact or proximal to the heart is anatomically classified as epicardial and pericardial adipose tissue^2,4^. Epicardial adipose tissue (eAT) lies between the myocardium and the visceral layer of the epicardium, whereas pericardial adipose tissue (pAT) is located further from the myocardium, between the visceral and parietal layers of the pericardium^4^. Both eAT and pAT depots are developmentally distinct in origin, with eAT being vascularized by coronary perfusion it has been linked to atrial fibrillation^5,6^, coronary artery disease^7^, and heart failure^8–10^. Pericardial fat has been associated with development of obesity-related CVDs, including coronary atherosclerosis^11^, heart failure^12^, vascular calcification^13^, and atrial fibrillation^14^.

Mitocondrial function is fundamental for the maintenance of adipose tissue homeostasis^15–17^. Obesity results in a reduction of AT mitochondrial content^18–20^, ATP production^21^, oxygen consumption^22^ alongside increased mitochondrial reactive oxygen species production (ROS)^23,24^ and fragmentation of this organelle^16^. These alterations contribute to metainflammation^15,25,26^. Moreover, brown adipocytes expressing uncoupling protein 1 (UCP1) dissipate energy as heat, increasing energy expenditure with the potential to combat metabolic diseases^27^. In mice and humans, brown adipose tissue activity status is positively associated with cardiometabolic health^28–30^, and an independent predictor of CVDs^31^. Although pAT exhibits certain characteristics of brown adipocytes, such as multilocular lipid droplets^32^, high metabolic rate^33^, and UCP1 expression^34^, its adaptation in obesity has not yet been fully investigated. Understanding this process may reveal therapeutic opportunities to combat obesity-related cardiometabolic diseases linked to this AT depot.

This study describes pAT presenting browning features in humans and mice. We show that pAT undergoes whitening in response to obesity in both species, while lipectomy of pAT in mice rescues some of the pathological cardiac changes. Finally, pharmacological activation of pAT by chronic administration of a β_3_-adrenergic agonist induced browning and improved heart function in obese mice in a pAT-dependent manner.

## Methods

### Ethics

All experiments were performed under institutional guidelines and approved by the Animal Care Committee at the University of Guelph under the protocol number 4142. All animal experiments were performed according to ARRIVE guidelines^71^ and NIH Guide for the Care and Use of Laboratory Animals.

### Human participants

Details regarding the design of the study are published elsewhere^72^. Briefly, all aspects of this study are in conformity to the Canadian Tri-Council Policy Statement on ethical conduct for research involving humans (TCPS-2–2014) and are in accordance with the World Medical Association Declaration of Helsinki^73^. The study has been registered with the National Clinical Trials Database of the NIH (www.clinicaltrials.gov).

All patients scheduled for elective, first-time cardiac surgery at the New Brunswick Heart Centre in Saint John, New Brunswick, and the Maritime Heart Centre in Halifax, Nova Scotia, were considered. During surgery, adipose tissue from subcutaneous, pericardial, and epicardial depots were collected in sterile specimen collection containers, labelled with a de-identification code and transferred to a research laboratory for analysis. The tissues will range in size from 0.5 to 1.5cm in width (0.3– 0.6cm thick) and were placed either in fixation buffer for histological analysis or liquid nitrogen for proteomic analysis.

### Mice

Male C57Bl/6N mice (12–18 weeks old) were randomly divided into two groups, a control group fed with control diet (10% energy from fat, Research Diets D12450J) and a HFD group fed with HFD (60% energy from fat, Research Diets D12492) for 8 weeks^24^. For the chronic CL-316,243 treatment, after 8 weeks of high-fat diet feeding, mice were randomized to receive either intraperitoneal (IP) injections of CL-316, 243 (0.2 mg/kg body mass; Sigma, C5976) or an equal volume of sterile saline (Vehicle) for 2 consecutive weeks, daily around 4 PM^61^. Body weight was assessed weekly. All mice were housed in the University of Guelph animal facility (22°C) on 12 h light–dark cycle with 24 h access to food and water *ad libitum*. Mice were anesthetized with an intraperitoneal injection of sodium pentobarbital (60 mg/kg, MTC Pharmaceuticals, Cambridge, ON) and tissues were harvested only after assurance of anaesthesia depth checked by leg retraction after toe pinch, palpebral reflex and movement of the whiskers. Once tissues were harvested, cervical dislocation was performed to euthanatize all mice.

### Pericardial adipose tissue removal

The thoracic open chest surgery was performed as previously published^44,74^. Briefly, mice were anesthetized with an isoflurane/oxygen mix (2%:100%), intubated and connected to a ventilator (Harvard Apparatus) and ventilated at 200 breaths per minute at 300 μL per breath. The 2^nd^ and 3^rd^ left ribs were separated from their cartilaginous connections with the sternum to expose the cardiac and pericardial adipose tissue. Pericardial adipose tissue was carefully excised using sharp scissor while preserving the cardiac tissue from damage. Sham surgery was similar to pAT removal, except ribs closed after separation without removing any pericardial adipose tissue.

### Echocardiography

Echocardiography of the LV was performed on anesthetized (isoflurane 2% induction followed by 0.5% maintenance) mice using Vevo2100 imaging systems (VisualSonics). B-mode and M-mode images of the LV were taken at the same time from the parasternal long axis of the heart^44^. Cardiac conductivity was assessed using pulsed-wave Doppler ultrasound and images were acquired in parasternal long axis/apical four-chamber view.

### Ex vivo insulin signaling

Ex vivo insulin signaling was performed as previously described with few modifications^75^. Pericardial adipose tissue was excised and weighed before being put in pre-warmed Media199 (Fisher Scientific, cat. No. 11043023) in the presence or absence of fresh diluted insulin (100nM NovoRapid®, Novo Nordisk). pWAT was incubated at 37°C under slow agitation for 15 min. Following this period, tissues were rapidly removed from the media, washed in PBS once, and flash-frozen in liquid nitrogen until further analysis to access Akt phosphorylation.

### Mitochondrial respiration

Mitochondrial respiration was determined in saponin-permeabilized adipose tissue in a respirometer chamber with 2 ml MiR05 at 37°C as previously described^24,50^. Briefly, adipose tissue was excised, immediately placed in BIOPS and minced with scissors and then transferred to MiR05 buffer. Tissue was weighed and rates of oxygen consumption by high-resolution respirometry (Oroboros Oxygraph-2k, Innsbruck, Austria) in the presence of saponin were determined. White adipose (iWAT and pWAT) mitochondrial respiration was tested in 15–20 mg of tissue with additions of substrates as follow: 5 mM pyruvate + 2 mM malate, 5 mM ADP, 5 mM glutamate, and 10 mM succinate; 25 μM dinitrophenol (DNP) was added to reach the maximal electron transport chain activity. Cytochrome *c* (10 μM) was used in all experiments to verify the integrity of the outer mitochondrial membrane.

### Mitochondrial reactive oxygen species production

Mitochondrial H_2_O_2_ emission was determined in left ventricle permeabilized cardiac fibers as previously described^44^. Briefly, fibers were inserted into a cuvette containing 10 μM Amplex Red (Invitrogen, Waltham, MA, USA), 40 U/mL SOD, and 5 U/mL horseradish peroxidase in Buffer Z at 25°C. Mitochondrial H_2_O_2_ emission rates were determined in the presence of 20 mM succinate in the presence of ADP 100 μM.

### Histology

Adipose tissue and a small portion of the anterolateral left ventricular tissue were fixed, embedded and stained with hematoxylin/eosin and picrosirius red, respectively, as previously described^24,50,44^. Images were taken using an Olympus FSX100 light microscope at a magnification of 40x. Fibrosis was expressed as a percentage of the total tissue area by averaging four different locations within the LV, whereas adipose tissue analysis is shown as adipocyte cross-sectional area averaging > 200 cells from 3 different images of the same animal/depot. Images were analysed using ImageJ 1.48 software (National Institutes of Health, Bethesda, MD, USA; https://imagej.nih.gov/ij/).

### Western blotting

Protein levels were determined by western blotting as previously described^24,75^. Samples were processed and loaded equally using SDS–PAGE gels, transferred to polyvinylidene difluoride membranes, and incubated in the appropriate blocking solution according to the antibody used. Antibodies used in this study include, total-Akt (1:1000, Cell Signaling Technology cat. no. 4691, Danvers, MA, U.S.A.), phosphorylated Akt-Ser473 (1:1000, Cell Signaling Technology cat. no. 9271), oxidative phosphorylation (mitochondrial content) cocktail (1:500, Abcam cat. no. Ab110413), UCP-1 (1:2000, Cell Signaling Technology cat. no. 10983), α-tubulin (1:5000, Abcam cat. no. ab7291, Eugene, OR, USA). All membranes were detected using enhanced chemiluminescence (ChemiGenius2 Bioimaging System, SynGene, Cambridge, U.K.) and densitometry was analyzed using ImageJ ((https://imagej.net/ij/).

### Protein Extraction and Purification for LC-MS

Protein extraction and bottom-up unlabeled proteomics were performed as previously described^76^. Twenty-five μg of protein per lysate was resolubilized with denaturation buffer (6 M urea/2 M thiourea), reduced with 10 mM DTT, and alkylated with 20 mM iodoacetamide at RT for 60 min. Samples were precipitated by adding 6:1 v/v cold acetone and kept at –80°C for 60 minutes and the pellet was collected after centrifugation at 10,000g for 10 min at 4°C. Samples were resuspended with 50 mM ammonium bicarbonate and MD-grade trypsin (Thermo Fisher Scientific, Cat#90057) was added overnight at 37°C. After digestion, samples were dried by vacuum centrifugation and purified using Pierce C18 Spin columns (Thermo Fisher Scientific, cat#89873).

The Vanquish Neo UHPLC system was coupled with Orbitrap Exploris 240 mass-spectrometer using the Easy-Spray source for nanoLC-MS protein identification. The mobile phase A and weak wash liquid was composed by water and 0.1% formic acid, and the mobile phase B and strong wash liquid was composed by 80% acetonitrile with 0.1% formic acid. The Orbitrap Exploris 240 MS was operated in data-dependent acquisition mode using a full scan with *m*/*z* range 375–1500.

### Proteomics data analysis

Analysis of the tissue proteome was performed in R (version 4.2.1) and R Studio (version 2024.04.2+764). Principal component analysis was performed using the Vegan package. P-values of differentially regulated proteins between groups were corrected as adjusted p-values by applying false discovery rate available in Deseq2 package^77^. Heatmaps and volcano plots were generated using all detected proteins or proteins sorted by p.adjust <0.1, gene ontology or MitoCarta3.0^36^. Gene set enrichment was performed with the Kyoto Encyclopedia of genes and genomes (KEGG)^78^ and EnrichR^79^.

### Statistical analysis

Results are expressed as individual observations with mean ± SD superimposed. Groups were compared using two-tailed Students t-test when comparing two groups or one-way ANOVA when three groups were compared. Categorical variables were analyzed using Fisher’s exact test. Outliers were removed when reached statistical significance by Grubbs’ test. Statistical analyses were performed using Prism 8.0 (GraphPad Software, La Jolla, CA, USA) or R (CRAN archive, version 4.3.3). Statistical significance was assumed when p<0.05 or adjusted p<0.1 for proteomics data. All codes used to analyze proteomic data are publicly available at: https://github.com/henver-brunetta

## Results

### Mitochondrial content is enriched in human pericardial adipose tissue

We started by investigating the human (N=3females/3males, age 61 ± 7 years, BMI 23 ± 1 kg/m^2^) adipose tissue proteome by applying bottom-up unlabeled mass spectrometry in three different adipose tissue depots: epicardial, pericardial, and subcutaneous adipose tissue (Fig. 1A). When comparing the absolute number of detected proteins in the three adipose depots from the same patient, we observed that eAT and pAT overlapped almost completely (Fig. 1B), sharing ca. 80-90% of the total amount of proteins identified. Although 1239 proteins were shared among the three depots, subcutaneous adipose tissue (scAT) shared much fewer proteins with the other two depots (1260 shared with pAT and 1668 shared with eAT), further highlighting the functional differences of this depot compared to the ones surrounding the heart. The 1239 conserved proteins among all depots were used for further principal component analysis (PCA) to confirm the proteome signature similarities between pAT and eAT, with almost completely overlapping ellipses and only 100 proteins differentially expressed between these two adipose depots (Supplementary Fig. 1A, B, Supplementary Table 1). In contrast, scAT showed a different pattern compared to pAT and eAT. Given transcriptional profile differences between eAT and scAT were already demonstrated ^4,35^, we next focused on the comparison between pAT and scAT for further analysis.

**Figure 1.**
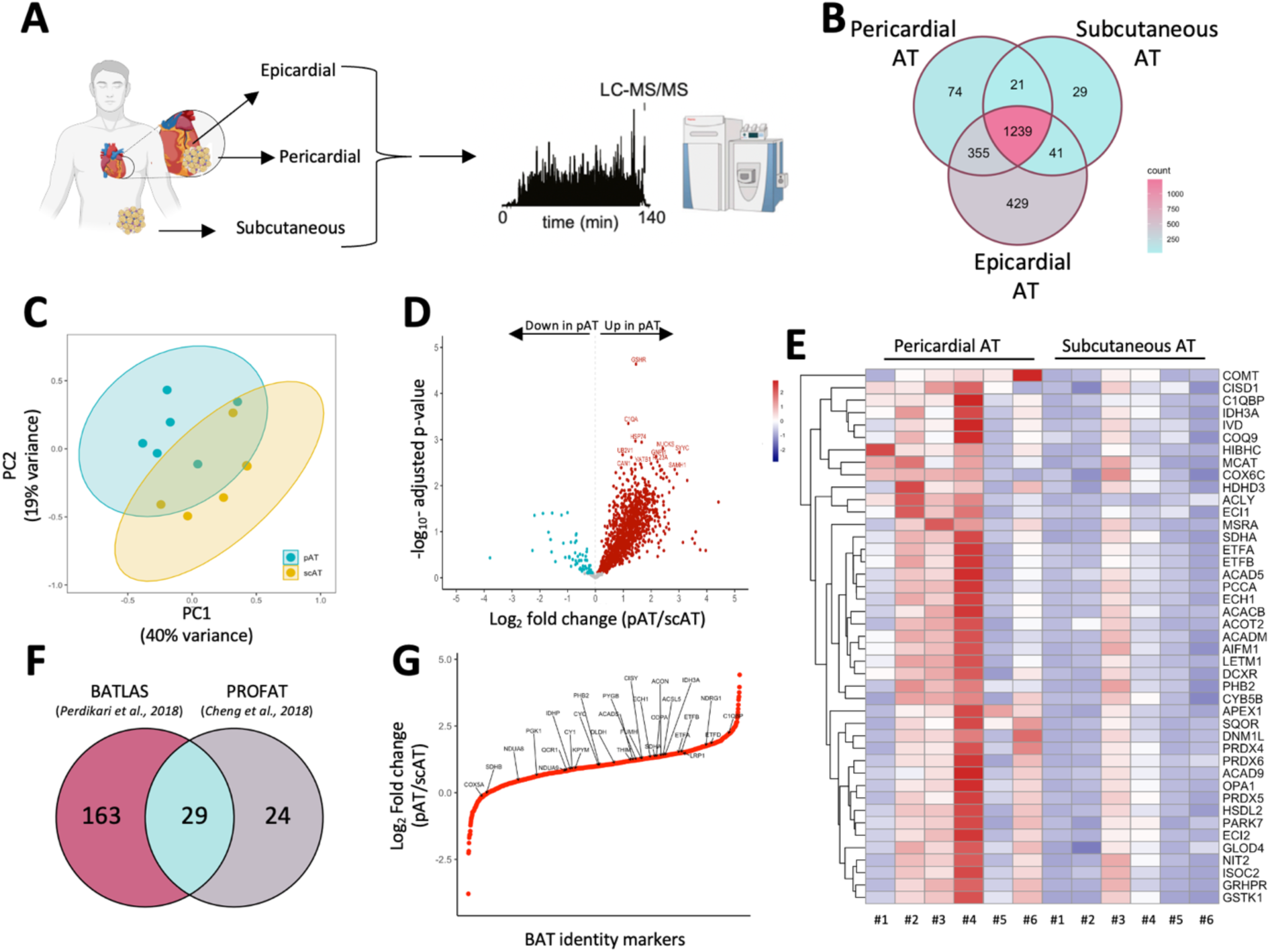
– Human pericardial adipose tissue presents browning and mitochondrial enrichment markers. (A) Schematic of anatomical location of adipose tissue biopsies from humans. (B) Venn diagram of identified proteins in pAT, eAT, and scAT (eAT, *n* = 6; pAT, *n* = 6, scAT, *n* = 6). (C) Principal component analysis and (D) volcano plot of the pericardial vs. subcutaneous adipose tissue (pAT, *n* = 6, scAT, *n* = 6). (E) Heatmap of differentially expressed mitochondrial proteins (pAT, *n* = 6, scAT, *n* = 6). (F) BAT identity markers in pAT and scAT selected from BATLAS and PROFAT and (G) enrichment of BAT identity markers in pAT compared to scAT (pAT, *n* = 6, scAT, *n* = 6). BAT – brown adipose tissue; eAT – epicardial adipose tissue; pAT – pericardial adipose tissue; scAT – subcutaneous adipose tissue; PC – principal component. Statistical test: two-tailed Student’s t-test. * adj.p-value < 0.1.

PCA of pAT compared to scAT demonstrated a clear proteome signature between these two depots (Fig. 1C). Volcano plot confirms these differences where almost 1/3 of the proteome is upregulated (adjusted p-value < 0.1) in pAT compared to scAT and a small fraction of proteins are less abundant in this depot compared to scAT (Fig. 1D, Supplementary Table 2). We also observed in 3 patients (50% of our sample) the presence of UCP1 in pAT (data not shown), while this protein was not detected in any of the scAT samples, suggesting that metabolic activity and mitochondrial bioenergetics could be linked to differences in the whole proteome of pAT. To test this, we used MitoCarta 3.0 ^36,37^ to filter mitochondrial proteins in our samples and noticed that among the differentially expressed proteins, almost all mitochondrial proteins were upregulated in pAT compared to scAT (Fig. 1E). We next calculated mitochondrial enrichment factor in our samples as a proxy of mitochondrial content by filtering out mitochondrial proteins from MitoCarta3.0 dataset and determining their sum intensities ratio to the whole proteome intensity. Supporting our previous observation of higher mitochondrial protein content in pAT compared to scAT, mitochondrial enrichment factor showed a strong trend towards enriched mitochondria in pAT (Supplementary Fig. 1C).

Given that mitochondrial content and the presence of UCP1^+^ cells are hallmarks of brown adipose tissue, we explored if other markers of brown adipose tissue would be found in pAT. To address this, we used BATLAS^38^ and PROFAT^39^, two independent algorithms based on human and mouse transcriptome to identify and predict brown adipocytes in mixed populations. Our proteome dataset detected 29 proteins confirmed to be BAT identity markers in both databases (Fig. 1F), which most of them (Fig. 1G), including mitochondrial (ACADS, C1QBP, COX5A, ETFA, IDH3A, SDHA), cholesterol (LRP1) and glucose (PYGB) metabolism and adipocyte differentiation proteins (NDRG1 and PHB2) ^40,41^, showed an enrichment in pAT compared to scAT. Altogether, human pAT exhibits a distinct proteome signature compared to subcutaneous AT, marked by an increased level of mitochondrial proteins, including the presence of UCP1 and BAT identity markers.

### Mouse pericardial adipose tissue presents a brown-like phenotype

Having mapped proteome pAT in humans, we next tested whether mice would present a similar profile. We analyzed pAT and scAT samples collected from adult male mice and quantified 1614 proteins in both depots, whereas each depot presented roughly ¼ of proteins that could not be detected in both (data not shown). Despite the majority of proteins being present in both depots, PCA depicted striking differences between pAT and scAT (Fig. 2B), while the volcano plot shows an almost equal number of proteins being up and downregulated in pAT compared to scAT in mice (Fig. 2C, Supplementary Table 3). KEGG analysis of the differentially expressed proteins shows enrichment of proteins involved in metabolism, such as PPAR signaling, fatty acid, and amino acid metabolism (Fig. 2D). Intriguingly, when proteins involved in lipid and glucose metabolism were plotted, we observed a downregulation in almost all proteins involved in lipid transport (i.e. FABP4, CPT2, CD36), lipid droplet formation (i.e. PLIN1, PLIN4), and fatty acid hydrolysis (i.e. HSL) in pAT compared to scAT whereas glucose handling proteins were overall upregulated in pAT (Fig. 2E). These data suggest that pAT is not only metabolically more active than scAT but also presents a shift in substrate handling capacity.

**Figure 2.**
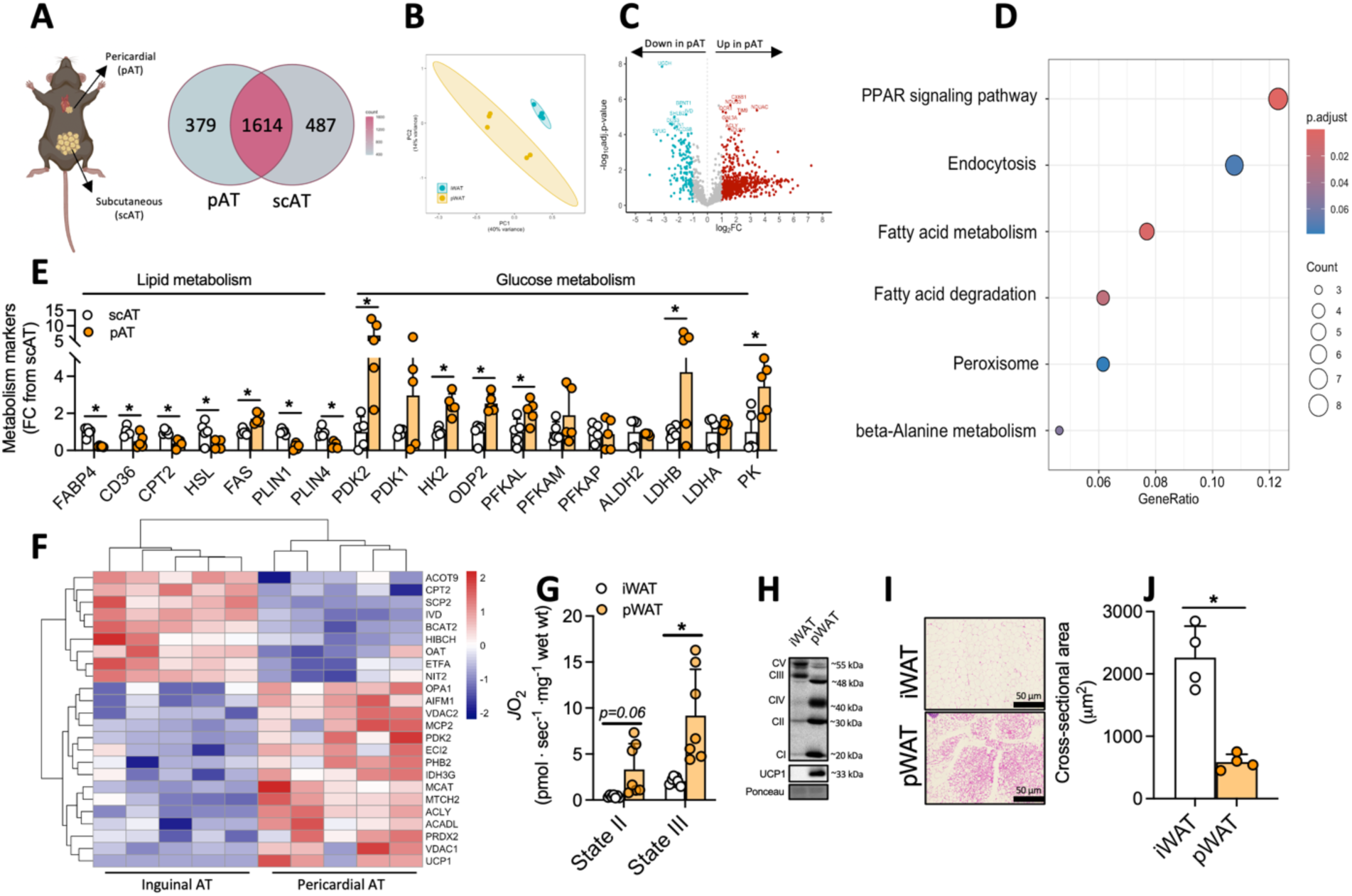
– Mouse pericardial adipose tissue presents a brown-like phenotype. (A) Venn diagram depicting conserved proteins found in pAT and scAT from mice. (B) Principal component analysis and (C) volcano plot comparing pAT and scAT proteome profile (scAT, *n* = 5; pAT, *n* = 5). (D) KEGG enrichment pathway analysis of the differentially expressed proteins between pAT and scAT. (E) Lipid and glucose metabolism related proteins in pAT vs. scAT. (F) Heatmap of differentially expressed mitochondrial proteins in pAT (scAT, *n* = 5; pAT, *n* = 5). (G) Mitochondrial O_2_ consumption in state II and state III in scAT and pAT (scAT, *n* = 6; pAT, *n* = 7). (H) Representative western blot highlighting the presence of UCP1 in pAT. (I) Representative and (J) quantification of cross-sectional area of histological images of scAT and pAT (scAT, *n* = 4; pAT, *n* = 4). CI – Complex I subunit NDUFB8; CII – Complex II subunit SDHB; CIII – Complex III subunit UQCRC2; CIV – Complex IV subunit COXII; CV – Complex V subunit ATP5A; pAT – pericardial adipose tissue; scAT – subcutaneous adipose tissue; PC – principal component. Statistical analysis: two-tailed Student’s t-test. * adj.p<0.1.

Given the mitochondrial proteome is upregulated in pAT from humans (Figure 1), we used MitoCarta 3.0 to identify mitochondrial proteins in our proteome dataset of differentially expressed proteins^36^. Interestingly, we observed the differences in the mitochondrial proteome are not uniform in pAT compared to scAT. While UCP1 is upregulated in pAT, several proteins involved in lipid oxidation were found to be downregulated (Fig. 2F). To test the functional differences between these two AT depots, we measured mitochondrial respiration in saponin-permeabilized pAT and scAT using pyruvate/malate (state 2) and ADP (state 3). We observed a strong trend towards higher respiration in pAT in state 2 and a significant elevated respiration in state 3 (Fig. 2G). Importantly, greater mitochondrial respiration was still present when glutamate and succinate were tested (Supplementary Fig. 2A), indicating the differences in O_2_ consumption between pAT and scAT are not substrate-specific, and likely due to greater mitochondrial content and UCP1 content (Fig. 2H). When histological analysis was performed in these two depots, we observed pAT adipocytes to be, on average, smaller compared to those from scAT (Fig. 2I, J) despite apparent heterogeneity of this depot (Supplementary Figure 2B). In summary, our data suggests that pAT presents features of brown adipose tissue, such as high expression of UCP1, mitochondrial respiration and small adipocytes.

### Obesity causes whitening of pericardial adipose tissue

During obesity, white adipose tissue expands considerably via adipocyte hypertrophy and hyperplasia, while brown adipose tissue presents markers of “whitening”. To investigate the effects of obesity on pAT adaptation, we fed male C57Bl/6N mice with a high-fat diet (60% calories from fat) for 8 weeks (Supplementary Fig. 3A). As expected, 8 weeks of HFD-feeding caused greater body weight gain and impaired glucose homeostasis (Supplementary Fig. 3B-D). Moreover, echocardiogram revealed a reduction in end systolic and end diastolic volume, cardiac output, and stroke volume (Fig. 3A-E) with smaller LV chamber diameter and no change in ejection fraction, fractional shortening and heart rate (Supplementary Fig. 3E-H). Moreover, Doppler analyses highlighted slower E and A waves (Supplementary Fig. 3I, J), and greater E/A ratios indicative of diastolic dysfunction (Fig. 3 F, G). In addition, although we found no differences in isovolumetric relaxation and contraction time, ejection time was shortened in HFD-fed mice (Supplementary Fig. 3K-M). In summary, HFD diet caused canonical hallmarks of cardiometabolic disease, including gain of adiposity, glucose intolerance, and cardiac dysfunction.

**Figure 3.**
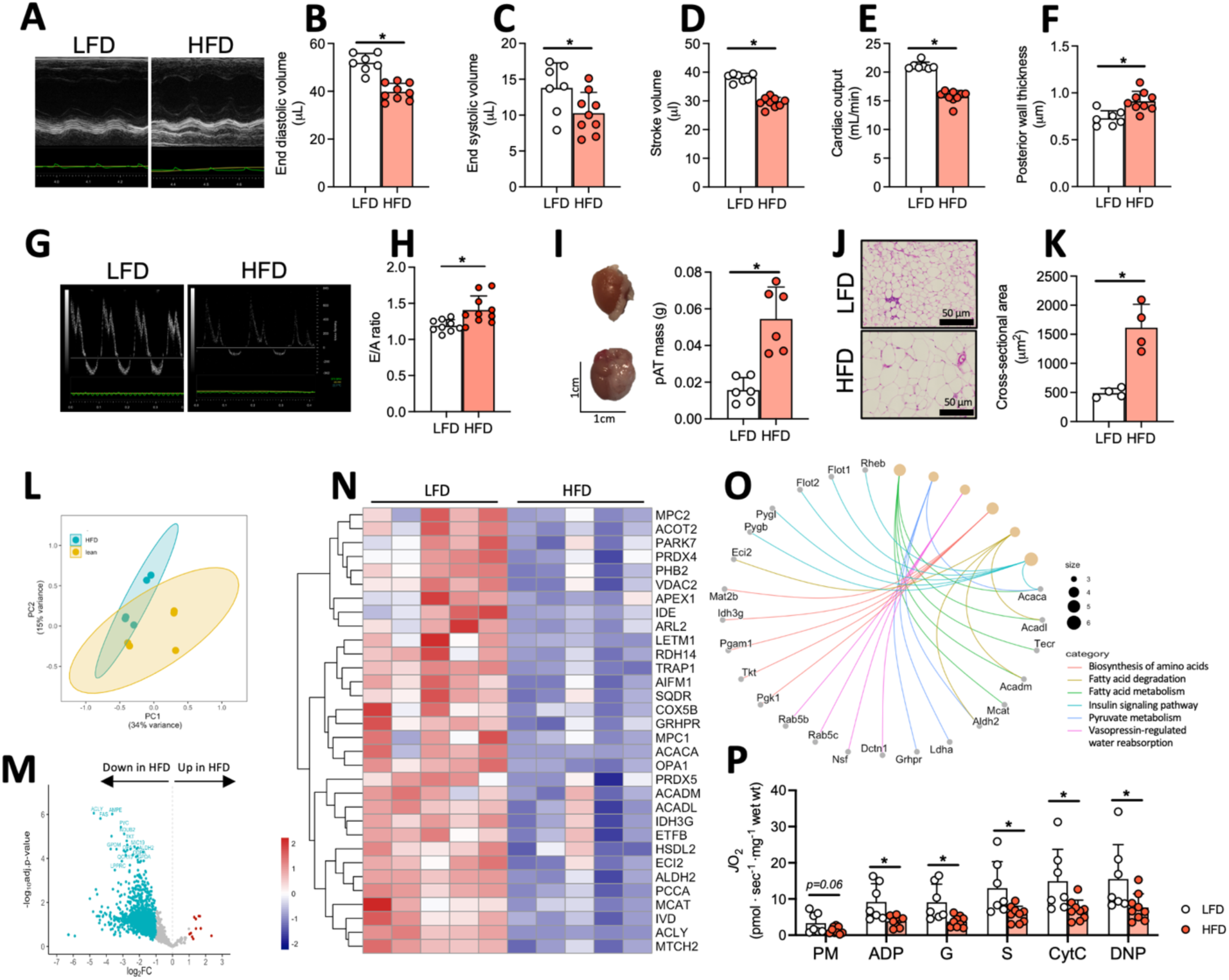
– High-fat diet feeding causes hypertrophy and whitening of pericardial adipose tissue. (A) Representative image of echocardiogram performed in low-fat and high-fat-fed mice. (B) End diastolic volume, (C) end systolic volume, (D) stroke volume, and (E) cardiac output (LFD, *n* = 7; HFD, *n* = 9). (F) Representative image of Doppler exam and (G) E/A ratio quantification (LFD, *n* = 9; HFD, *n* = 10). (H) pAT mass (LFD, *n* = 6; HFD, *n* = 6). (I) Histological representative images and (J) average cross-sectional area quantification of pAT from lean and obese mice (LFD, *n* = 4; HFD, *n* = 4). (K) Principal component analysis and (L) volcano plot of proteome of pAT from lean and obese mice (LFD, *n* = 5; HFD, *n* = 5). (M) Heatmap of the differentially expressed mitochondrial proteins from pAT of obese mice (LFD, *n* = 5; HFD, *n* = 5). (N) Gene concept network diagram of downregulated proteins in pAT. (O) Mitochondrial respiration assessed in pAT of both groups (LFD, *n* = 7; HFD, *n* = 9). pAT – pericardial adipose tissue; LFD – low-fat diet; HFD – high-fat diet; P – pyruvate; M – malate; ADP – adenosine diphosphate; G – glutamate; S – succinate; Cytc – cytochrome C; DNP – 2,4-dinitrophenol; PC – principal component. Scale bar = 50 μm. Statistical test: two-tailed Student’s t-test. * p-value < 0.05.

Next, we determined the effects of chronic HFD feeding on pAT. Pericardial adipose tissue presented whitening, as observed by heavier tissue mass (Fig. 3H) and enlarged adipocytes (Fig. 3I, J), similar to classical white adipose tissue depots. To further investigate the chronic effects of HFD on the pAT, we performed unlabelled proteomics in pAT harvested from low-fat and HFD-fed mice. PCA plot shows a clear separation between pAT from lean and obese mice (Fig. 3K), where the volcano plot demonstrates that a vast majority of proteins in pAT were downregulated in HFD-fed mice (Fig. 3L and Supplementary Table 4). KEGG analysis further demonstrates the enrichment of downregulated proteins found in pAT of HFD-fed mice on several key metabolic pathways of this tissue, such as fatty acid metabolism, biosynthesis of amino acids, pyruvate metabolism, fatty acid degradation and elongation, isoleucine degradation, and propanoate metabolism (Supplementary Fig. 3N). Given the mitochondrial enrichment found in pAT compared to scAT in lean animals (Fig. 2), we next filtered differently expressed proteins in our dataset using MitoCarta 3.0 to identify mitochondrial proteins affected specifically by HFD in pAT. Strikingly, all mitochondrial proteins differentially expressed in pAT from HFD-fed mice were downregulated (including a strong trend for UCP1, *adj.p-value=0.07*) compared to lean-derived pAT (Fig. 3M).

Gene-concept network diagram shows several proteins downregulated in pAT of HFD-fed mice were involved in insulin signaling, fatty acid and pyruvate metabolism (Fig. 3N), hallmarks of insulin resistance within adipose tissues. To investigate the functional effects of chronic HFD-feeding on mitochondrial function of pAT, we measured mitochondrial oxygen consumption over a wide range of substrates and energetic status. We observed an overall reduction in mitochondrial respiration, regardless of substrate offered or activation state of the electron transport chain (Fig. 3O), further confirming the functional consequences of reduced mitochondrial levels in pAT of HFD-fed mice. In summary, HFD-feeding induces whitening of pAT, characterized by adipocyte hypertrophy, low mitochondrial content and respiratory capacity, and insulin resistance.

### β-adrenergic agonist treatment induces browning in pericardial adipose tissue

Given the finding that whitening of pAT occurred with obesity-related cardiac dysfunction we next determined if browning of pAT could present a therapeutic target of opportunity. Using β-adrenergic agonists that are known to activate a browning program in adipocytes and shown some efficacy in cardiometabolic diseases^42^, has not been examined in the context of pAT before. To test this, we fed 12-15 weeks-old C57Bl/6N male mice a HFD for 8 weeks to induce obesity, glucose intolerance, and cardiac dysfunction (Supplementary Fig. 4A-M). At the end of the HFD-feeding, mice were randomly split into 2 groups: i) one receiving CL-316,243, a highly selective β3-adrenergic agonist, 0.2 mg/kg daily for 2 weeks and ii) another receiving volume-paired vehicle while animals were kept under HFD-feeding (Fig. 4A). No differences existed at time of randomization for body weight, fasting glucose, nor cardiac function between groups (Supplementary Fig. 5A-L). β3-adrenergic agonism slightly reduced body weight of HFD-fed mice improved whole-body glucose tolerance and induced browning in scAT (Supplementary Fig. 6A-H). Western blot of pAT revealed CL-316-243 treatment induced browning in this depot, as observed by marked increased in mitochondrial respiratory complexes subunits and the canonical brown adipose tissue marker UCP1 (Fig. 4B, C) in association with a marked reduction of pAT mass (Supplementary Fig. 6I). When mitochondrial O_2_ consumption was assessed using high-resolution respirometry, we observed an overall increase in mitochondrial respiration, including leak respiration and oxidative phosphorylation, further confirming the browning effect on mitochondrial biogenesis and uncoupling caused by β3-adrenergic agonist treatment (Fig. 4D). Combined, our experiments show that pAT is responsive to β3-adrenergic agonist treatment, increasing mitochondrial biogenesis and UCP1 expression even under chronic HFD-feeding.

**Figure 4.**
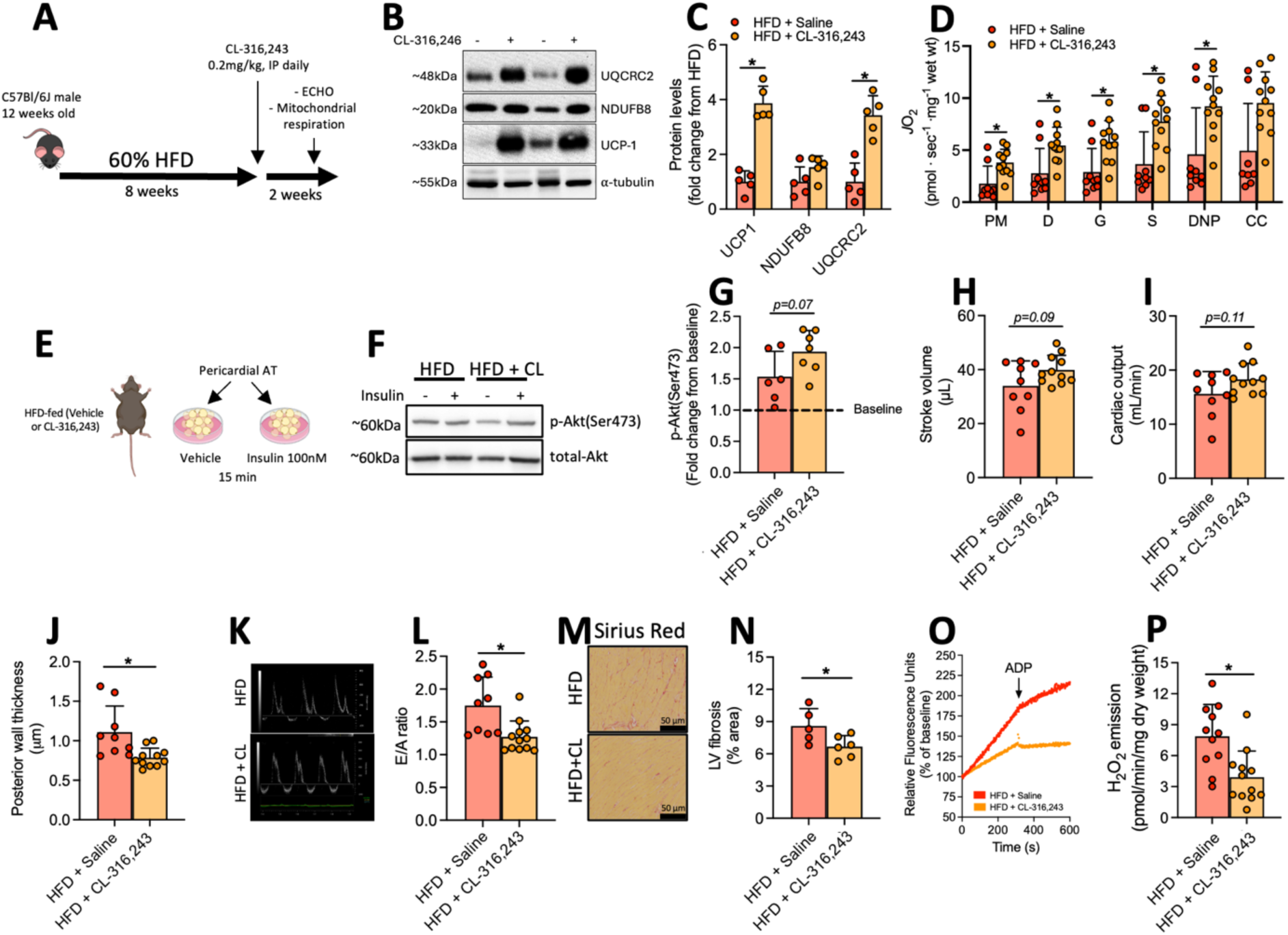
– β3-adrenergic agonism induces browning in pericardial adipose tissue. (A) Schematic representation of the experimental model of β3-agonist CL-316,243 treatment in HFD-fed mice. (B) Western blot images and (C) quantification of mitochondrial complexes subunits and UCP1 in pAT lysates (HFD + Saline, *n* = 5; HFD + CL-123,346, *n* = 5). (D) Mitochondrial respiration assessed in saponin-permeabilized pAT (HFD + Saline, *n* = 8; HFD + CL-123,346, *n* = 11). (E) Schematic model of *ex vivo* insulin stimulation in pAT and (F) representative western blot of total-Akt and Ser473-Akt phosphorylation levels. (G) Quantification of western blot experiments normalized by baseline p-Akt/total-Akt ratio (HFD + Saline, *n* = 6; HFD + CL-123,346, *n* = 7). (H) Stroke volume, (I) cardiac output, (J) and posterior wall thickness (HFD + Saline, *n* = 8; HFD + CL-123,346, *n* = 11). (K) Representative Doppler trace and (L) quantification of E/A ratio (HFD + Saline, *n* = 9; HFD + CL-123,346, *n* = 12). (M) Histological image of picrosirius red and (N) quantification of fibrotic area in left ventricle (HFD + Saline, *n* = 5; HFD + CL-123,346, *n* = 6). (O) Representative trace and (P) quantification of mitochondrial H_2_O_2_ emission in the presence of succinate and ADP in saponin-permeabilized LV fibers (HFD + Saline, *n* = 11; HFD + CL-123,346, *n* = 12). ADP – adenosine diphosphate; CL – CL-316-243; CC – cytochrome C; DNP – 2,4- dinitrophenol; G – glutamate; H_2_O_2_ - hydrogen peroxide; HFD – high-fat diet; *J*O_2_ – oxygen consumption; P – pyruvate; M – malate; NDUFB8 – Subunit of mitochondrial complex I; pAT – pericardial adipose tissue;S – succinate; UCP1 – Uncoupling protein 1; UQCRC2 – Subunit of mitochondrial complex III. Scale bar = 100 μm. Statistical test: two-tailed Student’s t-test. * p<0.05.

### Augmented browning of pericardial adipose tissue is associated with improvement in cardiac function

Brown adipose tissue activity is associated with cardiometabolic health^28^; therefore, we tested the effects of our β_3_-adrenergic agonist treatment on insulin signaling in pAT obtained from HFD-fed mice *ex vivo* (Fig. 4E). Though insulin had a mild effect on Akt-phosphorylation of pAT from HFD-fed mice, pAT from HFD-fed β_3_-adrenergic agonist-treated mice presented a trend (*p=0.07*) towards elevated Akt phosphorylation, averaging a ∼2-fold increase in the Akt-Ser473 phosphorylation levels (Fig. 4F, G). Given the strong relationship between adipocyte dysfunction and the development of cardiovascular diseases^21,22,43^, we also evaluated the β_3_-adrenergic agonism effect on cardiac function. An overall trend towards increased stroke volume (Fig. 4H, *p = 0.09*) and cardiac output (Fig. 4I, *p = 0.11*) in HFD-fed β_3_-adrenergic agonist-treated mice was observed. Moreover, β_3_-adrenergic agonism reduced posterior wall thickness and E/A ratio (Fig. 4J-L), previously augmented by obesity, indicating therapeutic efficacy.

Emerging evidence implicates fibrosis and mitochondrial redox imbalance as critical mediators of cardiac dysfunction in obesity ^44,45^. Markers of these underlying mechanisms related to cardiac structure and function with β3-adrenergic agonist in HFD- fed mice show reduced fibrotic area (Fig. 4M, N) and mitochondrial ROS emission in LV fibers (Fig. 4O, P). Altogether, browning of pAT by β3-adrenergic agonist treatment is associated with morphological and functional improvements in cardiac structure, stress, and function of HFD-fed mice.

### Pericardial adipose tissue accumulation contributes to cardiac dysfunction in obesity

The physiological role of pAT on the progression of cardiac dysfunction in obesity is unknown. Lipectomy of pAT did not affect overall function in lean animals (Supplementary Fig. 7A-J), suggesting a minor role of this depot on healthy mice. To determine if pAT contributes to the development of pAT to adapt to high-energy environments, we next subjected pAT-lipectomy and sham mice to HFD. Similar to LFD- feeding, the absence of pAT did not affect cardiac function through 6, 8, and 10 weeks after HFD-feeding (Supplementary Fig. 8A-J), ruling out a preventative role for pAT removal upon HFD-induced cardiac dysfunction.

Given that cardiovascular diseases in general occurs after the establishment of obesity, we then designed an additional experimental protocol aiming to address the therapeutic potential for pAT removal in mice with pre-existing obesity and cardiometabolic derangements. We fed 12–15-weeks-old C57Bl/6N male mice a HFD for 8 weeks to induce obesity and cardiac dysfunction, then performed either pAT lipectomy (no-pAT group) or sham surgery, keeping pAT intact (Sham group). After surgeries, animals continued to consume HFD for 3 weeks before re-assessing cardiac structure and function (Fig. 5A). Successful lipectomy was confirmed by significantly reduced pAT mass 3 weeks after the surgery showing little regeneration in the no-pAT group (Fig. 5B). While pAT removal did not affect end-systolic volume, end-diastolic volume, stroke volume, cardiac output or other markers of cardiac function (Fig. 5C-G and Supplementary Fig. 9A-C), we observed a reduction in posterior wall thickness of these animals compared to the Sham-group (Fig. 5H). Lipectomy had no difference in ejection time (Fig. 5L), E wave, A wave, or E/A ratio (Figure 5I, J and Supplementary Fig. 9D, E); however, we observed am increase in isovolumetric relaxation time (Fig. 5K) and a trend towards a decreased in isovolumetric contraction time (Fig. 5L, *p = 0.06*) with pAT removal. Mechanistically, pAT lipectomy with HFD did not affect fibrosis (Fig. 5M, N) nor mitochondrial ROS emission (Fig. 5O) in the LV, suggesting alternative mechanistic pathways beyond these common mediators of cardiac dysfunction. In sum, pAT lipectomy has no deleterious effects on cardiac function in lean animals nor prevents the development of dysfunction in obese mice. Yet, intervention pAT removal in established obesity can benefit posterior wall thickness and isovolumetric contraction/relaxation time. Taken together, the present data highlights potential for either pharmacological or surgical interventions targeting the pericardial adipose tissue for improved cardiac function in obese mice, suggesting expansion of pAT whitening is detrimental for cardiac function.

**Figure 5.**
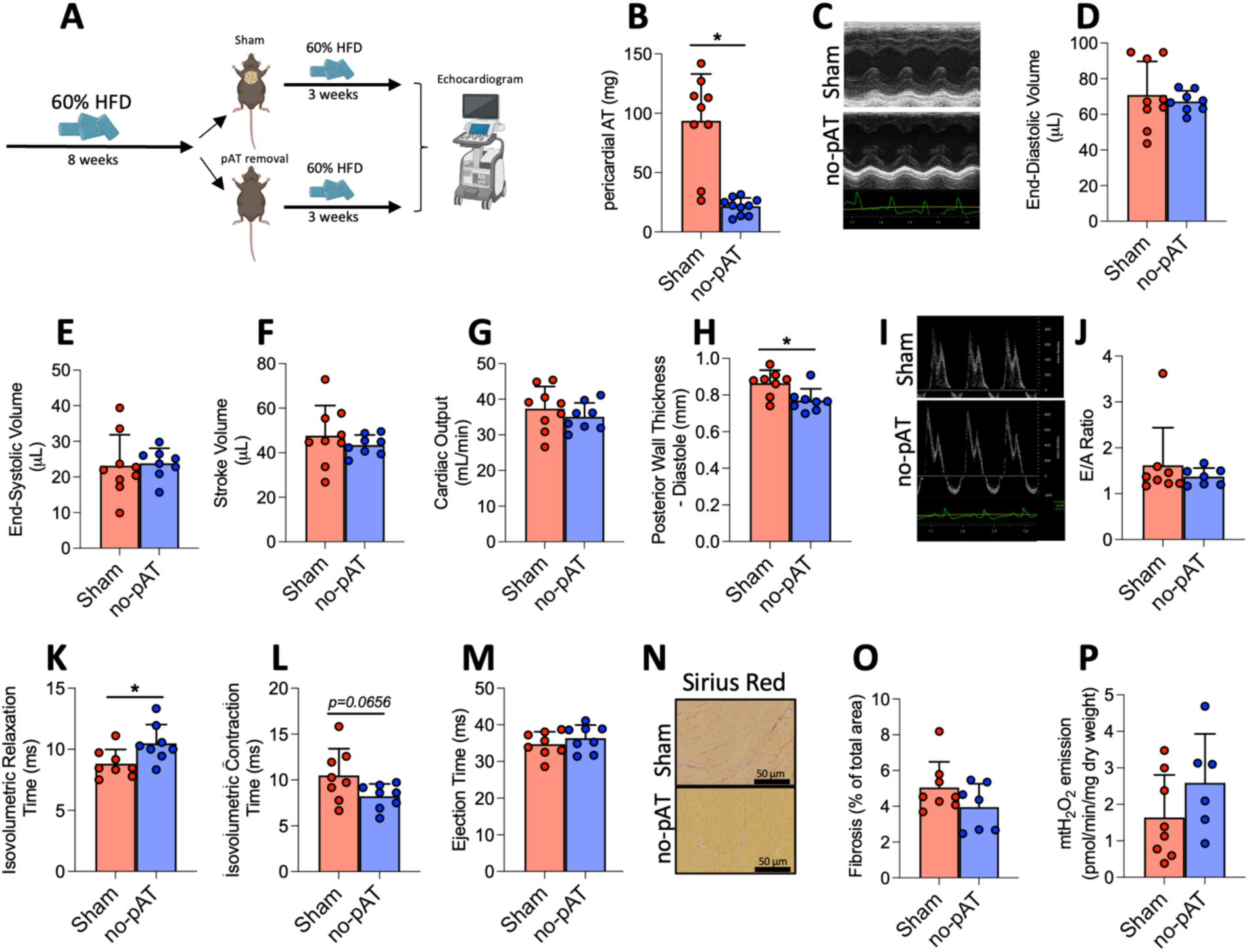
- Pericardial AT accumulation contributes to heart maladaptation in obesity. (A) Experimental design of surgical pAT removal. (B) Pericardial AT mass 3 weeks after lipectomy (Sham, n = 9; no-pAT, n = 10). (C) Representative images of echocardiogram. (D) End-diastolic volume, (E) end-systolic volume, (F) stroke volume, (G) cardiac output, (H) posterior well thickness during diastole (Sham, n = 9; no-pAT, n = 8). (I) representative image of Doppler ultrasound exam and quantification of (J) E/A ratio, (K) isovolumetric relaxation time, (L) isovolumetric contraction time, and (M) ejection time (Sham, n = 8; no-pAT, n = 8). (N) representative image of picrosirius red and (O) quantification of fibrotic area in the LF of HFD-fed mice (Sham, n = 8; no-pAT, n = 7). (P) Quantification of mitochondrial H_2_O_2_ emission in the presence of succinate + ADP (Sham, n = 8; no-pAT, n = 6). pAT – pericardial adipose tissue; LV - left ventricle; HFD – high-fat diet; mtH_2_O_2_ - mitochondrial hydrogen peroxide. Scale bar = 100 μm. Statistical analysis: two-tailed Student’s t-test. * p<0.05.

## β3-adrenergic agonism benefits depend on pericardial adipose tissue browning

Having observed that chronic β3-adrenergic agonism stimulates pAT browning and partially decreases indices of cardiac hypertrophy in HFD-fed mice (Figure 4), we next asked whether these physiological improvements would require pAT browning or an indirect effect of scAT browning or else simply the direct activation of β_3_-adrenergic receptors on myocardium^46,47^. To resolve this question and establish a causal role for pAT browning in cardiac function, we performed lipectomy surgery in chronic HFD-fed mice and let the animals recover for one week, while still under HFD-feeding, and then treated both groups daily with β_3_-adrenergic agonist as previously described for 14 days (Figure 6A). As expected, the pAT was smaller in the no pAT group compared to the Sham group after β_3_-adrenergic agonist treatment (Fig. 6B). Despite β_3_-adrenergic agonist treatment, we identified that the improvements in stroke volume and cardiac output were abrogated in the absence of pAT in obese mice (Fig. 6C-E).

**Figure 6.**
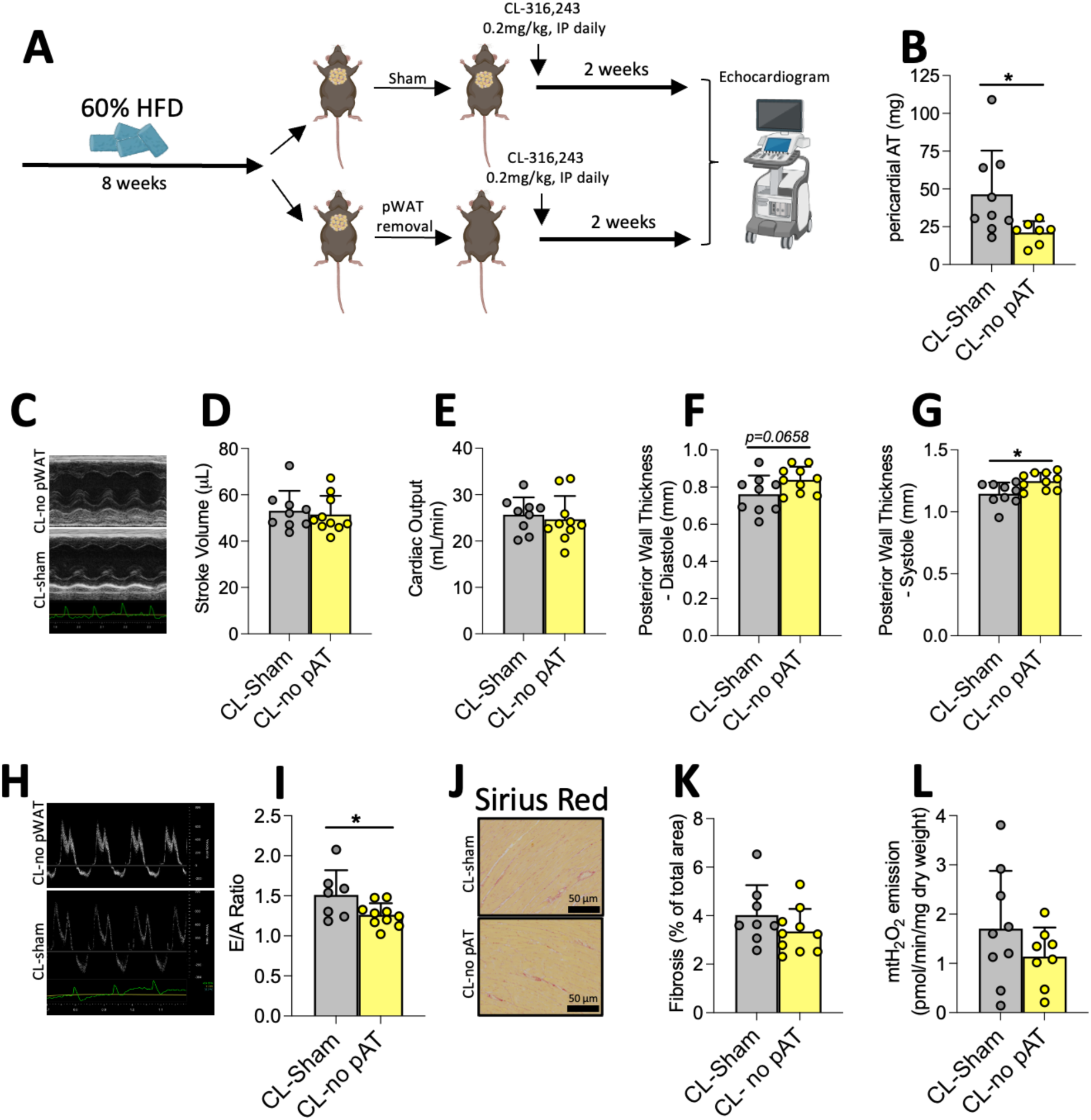
Cardiac benefits from β3-adrenergic agonism are dependent on pAT browning. (A) Experimental design of pAT removal from HFD-fed mice following by chronic CL-316,243 treatment. (B) Pericardial adipose tissue mass following lipectomy (CL-Sham, *n* = 9; CL-no pAT, *n* = 7). (C) Representative image in M-mode of echocardiographic exam. Quantification of (D) stroke volume, (E) cardiac output, (F) posterior wall thickness during diastole and (G) systole (CL- Sham, *n* = 9; CL-no pAT, *n* = 10). (H) Representative image of Doppler ultrasound exam and (I) E/A ratio (CL-Sham, *n* = 7; CL-no pAT, *n* = 10). (J) Representative image of picrosirius red and (K) quantification of fibrotic area in the LV of HFD-fed mice (CL-Sham, *n* = 8; CL-no pAT, *n* = 10). (L) Quantification of mitochondrial H_2_O_2_ emission in the presence of succinate + ADP (CL-Sham, *n* = 9; CL-no pAT, *n* = 8). pAT – pericardial adipose tissue; HFD – high-fat diet. Scale bar = 100 μm. Statistical analysis: two-tailed Student’s t-test. * p<0.05.

Moreover, the absence of pAT not only eliminated the beneficial effect on LV wall hypertrophy but exacerbated wall thickening in β_3_-adrenergic agonist-treated obese mice (Fig. 6F, G). This paradoxical worsening provides particularly strong evidence that pAT is the primary mediator of the beneficial effects cardiac function, an improvement in β_3_- adrenergic agonist treatment. Interestingly, E/A ratio was improvements were maintained despite pAT removal (Fig. 6H, I), suggesting some independent effects of β_3_-adrenergic agonist treatment in cardiomyocytes. Finally, the beneficial effects of β_3_-adrenergic agonist treatment on LV fibrosis area and mitochondrial ROS emission were abrogated in the absence of pAT in HFD-fed mice (Fig. 6J-K). In summary, the induction of pAT browning by β_3_-adrenergic agonism is necessary for the amelioration of major cardiac dysfunction in obese mice.

### Obesity affects the mitochondrial proteome of human pericardial adipose tissue

Finally, we extended our findings to a clinically relevant experimental setting where we collected pAT samples from patients with a BMI < 25 kg/m^2^ (Lean) and greater than 30 kg/m^2^ (Obese) and performed unlabelled proteomics in these samples (Fig. 7A). To confirm our cohort’s phenotype, BMI was significantly higher in obese patients associated with greater waist-to-hip ratio (Table 1). In addition to the expected anthropometric differences, we also observed higher fasting blood glucose and type 2 Diabetes prevalence, lower LDL cholesterol and HDL levels in the obese group, hallmarks of metabolic complications caused by obesity. Regarding cardiac function, obese patients performed ca. 40% worse in the 6-minutes maximal walking test compared to the lean patients.

**Figure 7.**
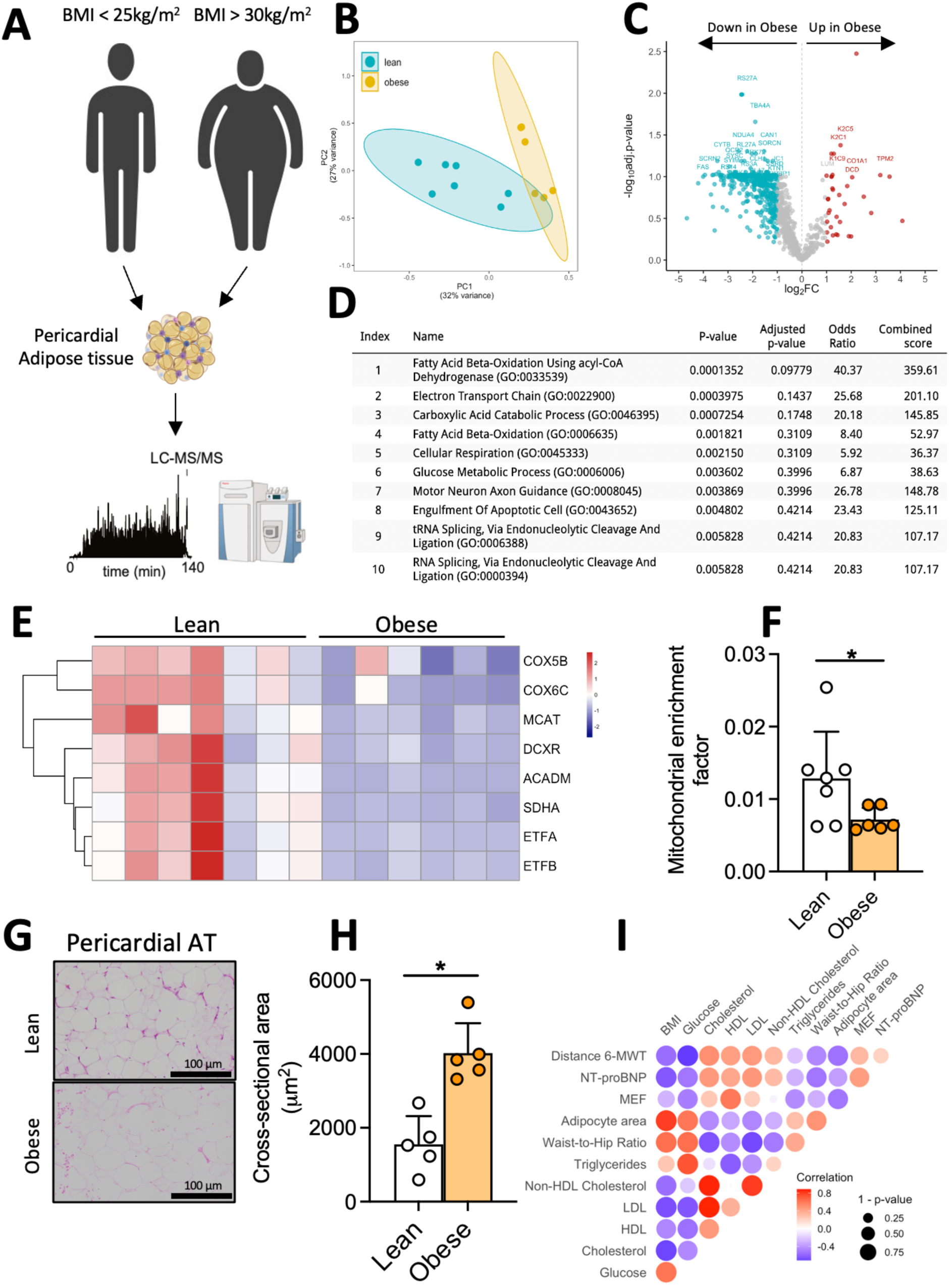
– Obesity negatively impacts mitochondrial proteome of pericardial adipose tissue in humans. (A) Workflow of proteomics of pericardial adipose tissue obtained from lean (BMI < 25 kg/m^2^) and obese (BMI > 30 kg/m^2^) patients. (B) Principal component analysis and (C) volcano plot of proteome from pAT of lean and obese patients (Lean, *n* = 7; Obese, *n* = 6). (D) Enrichment analysis of the downregulated proteins in the pAT of obese patients. (E) Heatmap of proteins present in the six most enriched pathways obtained from downregulated proteins found in the pAT of obese population (Lean, *n* = 7; Obese, *n* = 6). (F) Mitochondrial enrichment factor calculated from proteome dataset (Lean, *n* = 7; Obese, *n* = 6). (G) Histological images (scale bar = 100μm) and (H) quantification of adipocyte area from pAT and scAT of humans with and without obesity (Lean, *n* = 5; Obese, *n* = 5). (I) Correlation analysis of MEF in function of clinical parameters of patients, r values are color coded and only p<0.05 circles are shown. BMI – body mass index; MEF – mitochondrial enrichment factor; NT-proBNP - N terminal-pro brain natriuretic peptide; pAT – pericardial adipose tissue; scAT – subcutaneous adipose tissue; PC – principal component. Statistical analysis: two-tailed Student’s t-test. * p<0.05.

**Table 1.** – Clinical parameters of patients.

|  | Lean (*n* = 7) | Obese (*n* = 6) | *p*-value |
| --- | --- | --- | --- |
| Age (years) | 61.28 ± 7.04 | 60.50 ± 7.39 | 0.8481 |
| BMI (kg/m<sup>2</sup>) | 23.05 ± 1.63 | 34.03 ± 5.5 | 0.0004 |
| Fasting glucose (mmol/L) | 4.89 ± 0.58 | 9.52 ± 2.08 | 0.0001 |
| Cholesterol (mmol/L) | 4.28 ± 0.65 | 3.05 ± 0.52 | 0.0034 |
| HDL (mmol/L) | 1.52 ± 0.35 | 0.96 ± 0.29 | 0.0098 |
| LDL (mmol/L) | 2.31 ± 0.68 | 1.07 ± 0.33 | 0.0040 |
| Non-HDL Cholesterol (mmol/L) | 2.78 ± 0.83 | 2.09 ± 0.51 | 0.1078 |
| Triglycerides (mmol/L) | 1.04 ± 0.47 | 2.32 ± 1.42 | 0.0454 |
| Waist-to-hip ratio | 0.87 ± 0.07 | 1.02 ± 0.07 | 0.0052 |
| Urea (mmol/L) | 6.16 ± 1.96 | 6.20 ± 2.05 | 0.6064 |
| WBC (x10<sup>9</sup>/L) | 6.58 ± 0.87 | 7.36 ± 1.97 | 0.5824 |
| RBC (x10<sup>12</sup>/L) | 4.44 ± 0.42 | 4.65 ± 0.47 | 0.4603 |
| Creatinine (μmol/L) | 87.6 ± 19.7 | 76.8 ± 19.8 | 0.5804 |
| Glycated hemoglobin (mmol/L) | 5.96 ± 1.37 | 6.88 ± 0.94 | 0.2622 |
| 6-MWT (meters) | 384 ± 62 | 236 ± 81 | 0.0036 |
| Diabetes | (1/7) | (5/6) | 0.0291 |

BMI – body mass index; HDL – high-density lipoprotein; LDL – low-density lipoprotein. WBC - white blood cells; RBC - red blood cells; 6-MWT - 6 minutes maximal walking test. Statistical test: Two-tailed Student's t-test or Fisher exact test for categorical variables. \* *p*<0.05.

In the proteomics of pAT of these patients, PCA performed on the common proteins detected in all patients shows a distinct proteome signature of pAT samples obtained from obese compared to lean donors (Fig. 7B), indicating that obesity modulates the pAT proteome signature in humans. Moreover, similar to what we found in mice, the vast majority of proteins identified in pAT were downregulated in obese patients, whereas only a dozen proteins were upregulated in this population compared to lean patients (Fig. 7C, Supplementary Table 5). Enrichment analysis of the downregulated proteins in pAT from obese patients shows several metabolic pathways downregulated in this depot, such as fatty acid β-oxidation, electron transport chain, carboxylic acid catabolic process, cellular respiration, and glucose handling (Fig. 7D), suggesting a whitening of pAT in humans caused by obesity. By using gene ontology from the 6 most enriched pathways (i.e. GO:0033539, GO:0022900, GO:0046395, GO:0006635, GO:0045333, GO:0006006), we created a heatmap of the proteins identified in our dataset belonging to these pathways. As expected, all proteins found in the enrichment analysis were downregulated in the pAT of obese patients (Fig. 7E), with several belonging to mitochondrial oxidative phosphorylation processes.

We next calculated the mitochondrial enrichment factor (MEF) in these samples using MitoCarta3.0. Indeed, MEF was reduced in obese pAT compared to lean patients (Fig. 7F). Moreover, as observed in mice, obesity caused hypertrophy of adipocytes in pAT (Fig. 7G, H) to the same degree it caused in scAT (Supplementary Fig. 10A, B). Interestingly, although BMI was positively correlated with adipocyte size, fasting blood glucose levels, and waist-to-hip ratio, we did not observe a significant correlation of MEF and BMI (Fig. 7I). However, MEF was negatively correlated with adipocyte size (r=-0.52) and fasting blood glucose levels (r=-0.49) (Fig. 7I), further suggesting an impairment in mitochondrial metabolism as a function of adipocyte size in pAT and whole-body glucose homeostasis. In order to verify the role of pAT in the development of obesity associated cardiovascular disease, we assessed correlation of MEF within pAT with fitness performance in our patients through the 6 minutes maximal walk distance test (6-MWT). Interestingly, MEF was positively associated with 6-MWT (r=0.32), reinforcing the potential implications of mitochondrial content in pAT to the cardiovascular function. We also determined the circulating levels of NT-pro BNP, a marker of cardiac remodelling associated with heart failure. Interestingly, while we observed a negative correlation between BMI and waist-to-hip ratio with NT-pro BNP (r=-0.64) in our cohort (Figure 7I), supporting previous reports ^48,49^, there was a positive correlation (r = 0.44) between NT-pro BNP levels and MEF within the pAT, supporting the finding that obesity and adipocyte hypertrophy affect MEF and NT-pro BNP^41,42^. In summary, obesity causes pAT whitening in humans, characterized by greater adipocyte area, with diminished mitochondrial content and oxidative capacity, parallel to those seen in mice.

## Discussion

Here, we provide a mechanistic dissection of pAT plasticity and its relationship with obesity-induced cardiac dysfunction. We found in mice and humans that pAT presents a browning signature, as indicated by the enrichment of mitochondria, UCP1 expression, and greater levels of BAT identity markers. Moreover, we observed whitening of pericardial AT in obesity, characterized by adipocyte hypertrophy, mitochondrial content loss, and decreased UCP1 levels, suggesting a maladaptive response that may contribute to obesity-related cardiac dysfunction. Pharmacological browning and lipectomy of pAT improved some of the pathological adaptative responses in the heart of obese mice. Lastly, the thermogenic potential of pAT is required for β_3_-adrenergic agonist mediated improvements in heart function.

While adipose function varies according to the depot studied^3,17,50^, the most documented differences are between subcutaneous and visceral adipose tissue, where accumulation of visceral adipose tissue presents a higher risk for the development of CVDs^51,52^. Regarding adipose tissue surrounding the cardiac tissue, eAT has been associated with a wide range of cardiac diseases^4,6,8,9,35,53,54^. We found a similar proteome signature between eAT and pAT, suggesting despite their different ontogenesis the two depots have comparably consistent proteome profiles. Proteome signature of pAT compared to subcutaneous suggests a more metabolically active tissue, given that several BAT identity markers were found to be differentially expressed in pAT compared to scAT^38^. Despite such intrinsic differences, pAT was not shielded from a high caloric environment, given we found a similar pAT plasticity as commonly observed in subcutaneous white adipose tissue^17,55^. Confirming the maladaptive role of pAT in obesity, lipectomy of this depot partially rescued the obese cardiac phenotype in mice. Indeed, a case study of an obese diabetic human revealed that lipectomy of excessive mediastinal AT, including pAT, improved compression of the right heart following complications from a triple coronary artery bypass surgery^56^. This case suggests lipectomy could be a prophylactic approach in open heart surgeries, though systematic studies should be performed to confirm any beneficial effects of lipectomy of any adipose depot in obese patients.

Activation of brown adipose tissue holds promising therapeutical potential to counteract the deleterious effects of chronic positive energy balance^57–59^. However, despite such enthusiasm, the low amount of brown adipose tissue in adults challenges the efficacy of such interventions^60^. As an alternative, it has been suggested that white adipocytes could acquire BAT-like features, such as UCP1 expression and thermogenic potential^61,62^. In HFD-fed mice, β3-adrenergic agonism as form of treatment increased thermogenic potential, strongly suggesting the browning capacity of the pAT depot even under a high-caloric environment. Importantly, increased browning status of pAT following β3-adrenergic agonism was associated with improved cardiac function, LV fibrosis and thickness, reinforcing the potential therapeutical capacity that browning of pAT can have on cardiac function^29,30,58^. The blunted beneficial effects of β_3_-adrenergic treatment following lipectomy of this tissue suggest paracrine communication between pAT and the heart that is independent of systemic effects of subcutaneous adipose browning. Beyond its thermogenic potential, BAT also secretes a wide range of microRNAs, proteins, and lipids with endocrine action^3,63–65^. Therefore, it is plausible to speculate the improvements in heart function following β_3_ agonist treatment could be a result of local altered secretory pattern from pAT, in addition to its increased thermogenic activity. Nevertheless, this hypothesis remains to be tested.

Although we observed a negative correlation between MEF and both adipocyte size and fasting blood glucose levels, no significant correlation with BMI could be a result of BMI’s recognized limitations^66,67^. Alternatively, adipocyte area is considered a predictor of adipose tissue dysfunction and metabolic disarrangement^68^. Lower mitochondrial content has been observed in previous studies in visceral and subcutaneous depots^18,21,22,43,69^. Our data point to a negative correlation between mitochondrial content, pAT function, and overall health in both mice and humans. While pharmacological approaches that target mitochondrial content in adipocytes have been shown to improve overall health in an obese population^70^, whether these approaches can also target pAT will require future exploration.

Taken together, we found a distinct proteome signature between pAT and other AT depots in humans and mice. In addition, we demonstrated the whitening of pAT in obesity contexts in both species. Finally, we demonstrated the therapeutical potential that pAT holds to cardiometabolic diseases by responding to chronic β_3_-adrenergic treatment and lipectomy. Given the importance and plasticity of this depot, we conclude that the browning status of pAT regulates cardiac function in obesity-related preclinical and clinical contexts.

## Acknowledgements

We thank Dr. Diana Philbrick, Dr. Dyanne Brewer for their technical assistance in this project. Human tissues and clinical data were collected and collated in partnership with the IMPART BioBank (https://impart.team/biobank/). The authors extend their appreciation to the patient participants, the technical support of Saumil Shah, Dr. Christie Aguiar, and Hany Motawea, and for financial support of the Dalhousie Medical Research Foundation, Heart & Stroke Foundation of New Brunswick, and Saint John Regional Hospital Foundation - Chesley Family.

---

**Supplementary Figure 1.**
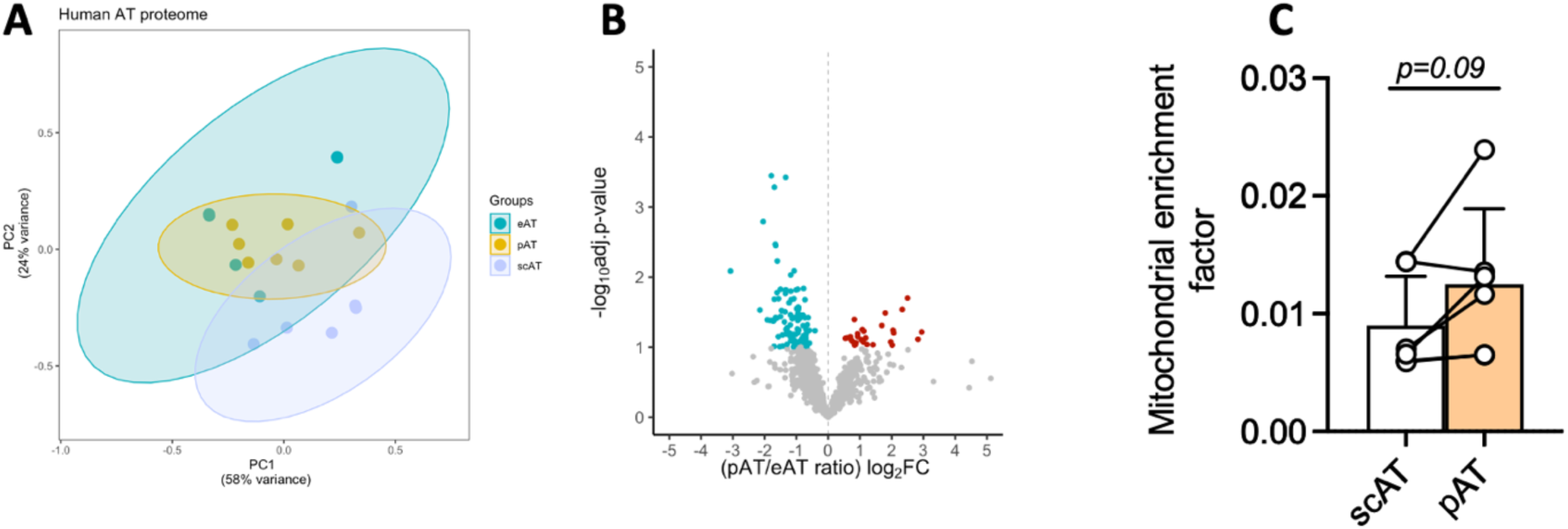
- Proteome of human adipose tissue. (A) Principal component analysis of all conserved detected proteins of subcutaneous, epicardial, and pericardial adipose tissue. (B) Volcano plot comparing pericardial and epicardial human adipose tissue. (C) Mitochondrial enrichment factor in subcutaneous and pericardial human adipose tissue. PC - principal component; pAT - pericardial adipose tissue; eAT - epicardial adipose tissue; scAT - subcutaneous adipose tissue.

**Supplementary Figure 2.**
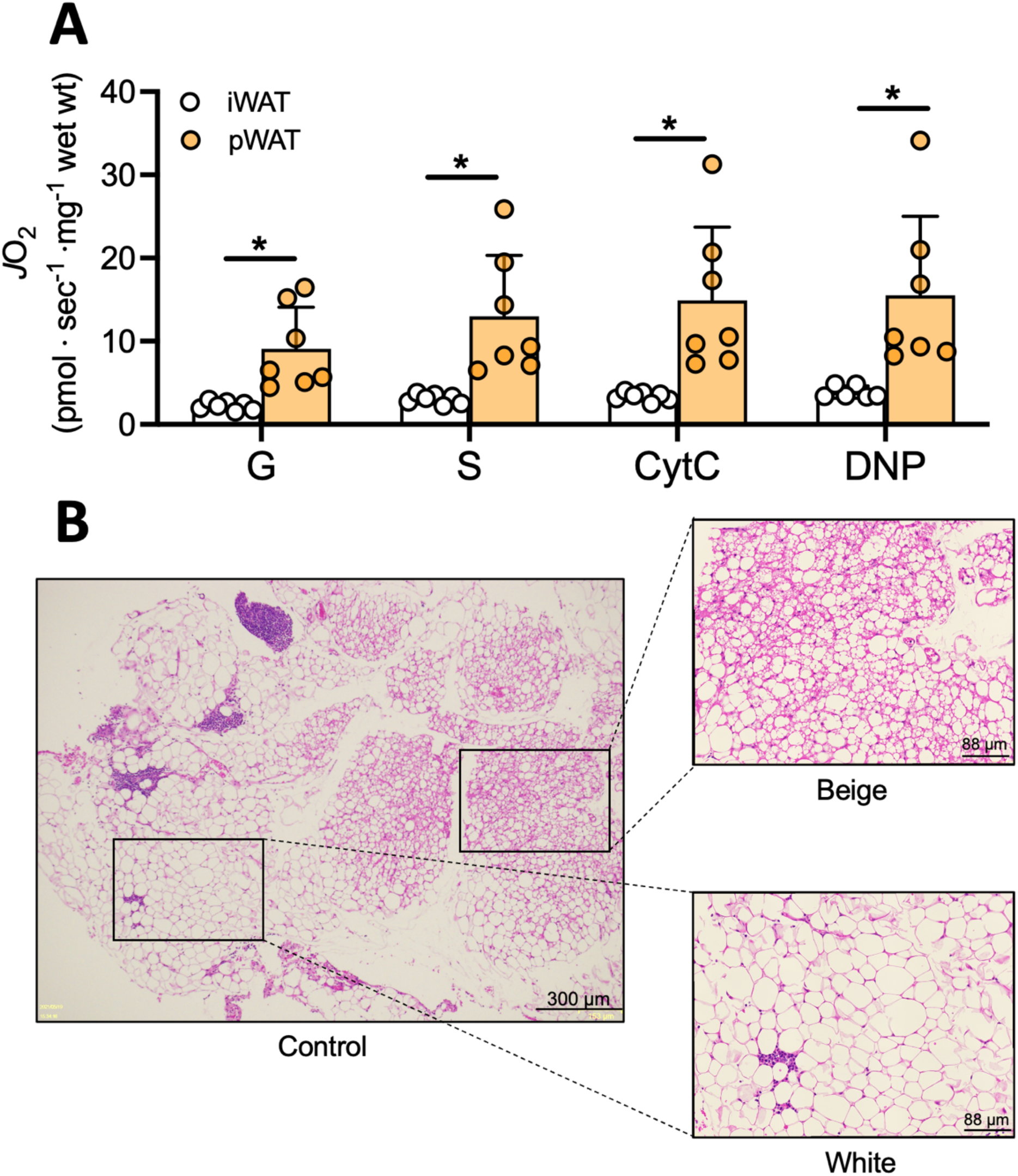
– Mitochondrial respiration and pericardial AT heterogeneity. (A) Mitochondrial respiration in the presence of ADP stimulated by succinate, glutamate, cytochrome c, or DNP. (B) Pericardial adipose tissue heterogeneity. G – glutamate; S – succinate; CytC – cytochrome c; DNP - 2,4-Dinitrophenol. Statistical analysis: two-tailed Student’s t-test. * p<0.05.

**Supplementary Figure 3.**
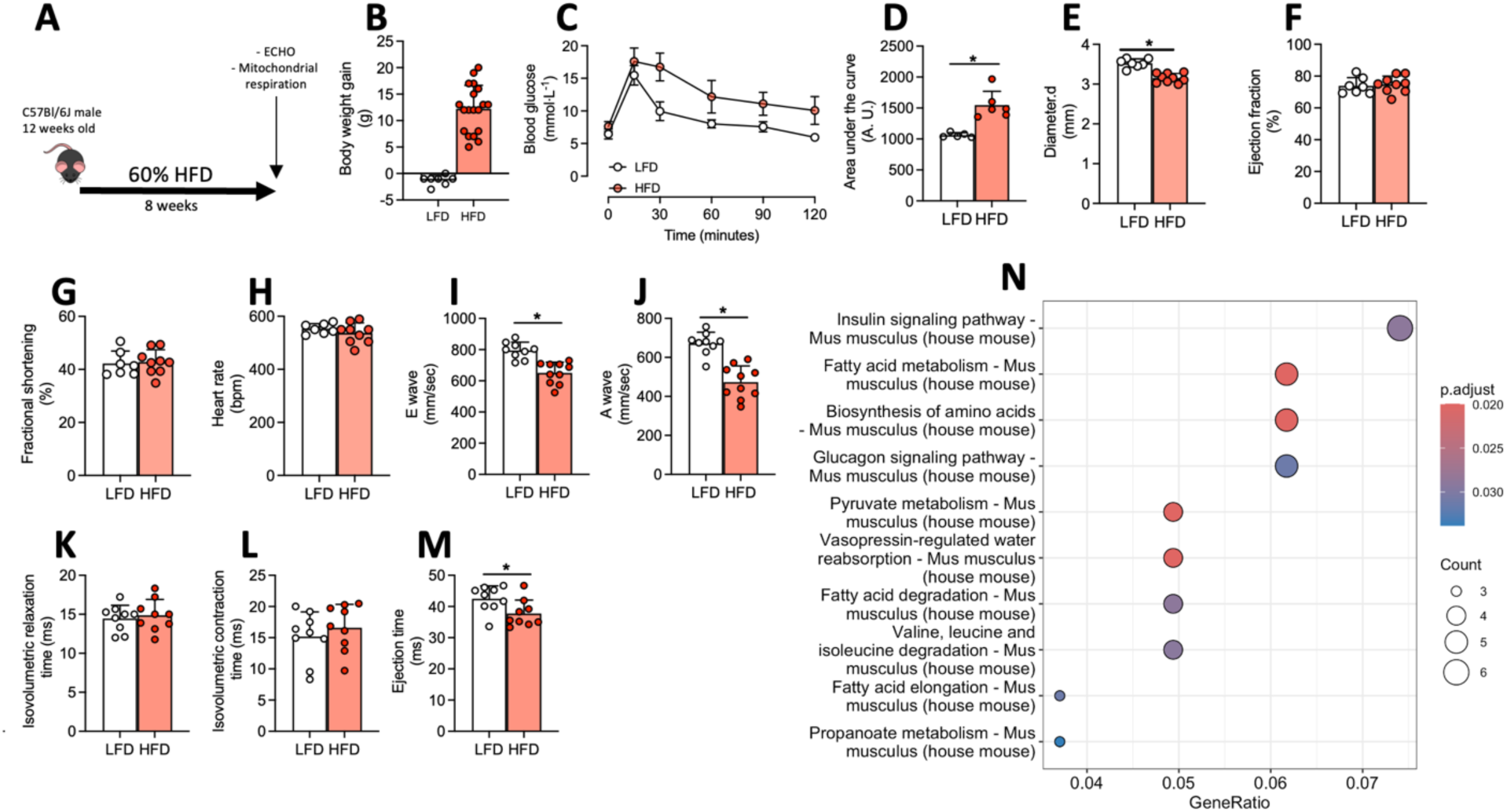
– Chronic HFD feeding induces obesity, glucose intolerance, and heart dysfunction. (A) Experimental design of 8 weeks of HFD-feeding. (B) Body weight gain, (C) glucose excursion during ipGTT and (D) area under the curve. (E) LV diameter during diastole, (F) ejection fraction, (G) fractional shortening, (H) heart rate, (I) E’ wave, (J) A wave, (K) isovolumetric relaxation time, (L) isovolumetric contraction time, and (M) ejection time. (N) KEGG enrichment analysis. ECHO – echocardiogram exam; LFD – low-fat diet; HFD – high-fat diet. Statistical analysis: two-tailed Student’s t-test. * p<0.05.

**Supplementary Figure 4.**
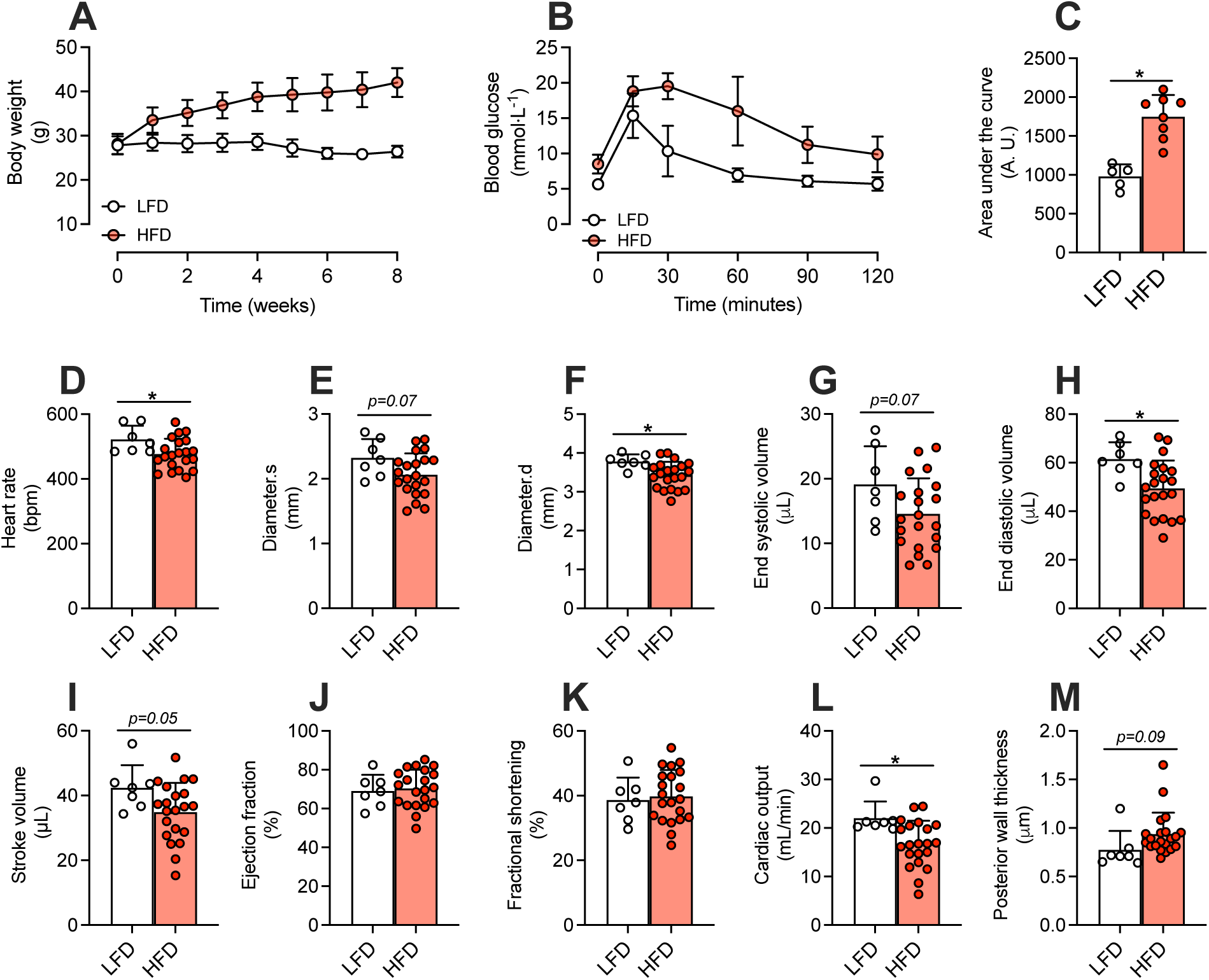
– Cardiometabolic effects of HFD before starting CL- 123,346 treatment. (A) Body weight during 8 weeks of HFD, (B) glucose excursion during ipGTT and (C) area under the curve. (D) heart rate, (E) LV diameter during systole and (F) diastole, (F) ejection fraction, (G) end systolic volume, (H) end diastolic volume, (I) stroke volume, (J) ejection fraction, (K) fractional shortening, (L) cardiac output, and (M) posterior wall thickness. HFD – high-fat diet; LFD – low-fat diet. Statistical analysis: two-tailed Student’s t-test. * p<0.05.

**Supplementary Figure 5.**
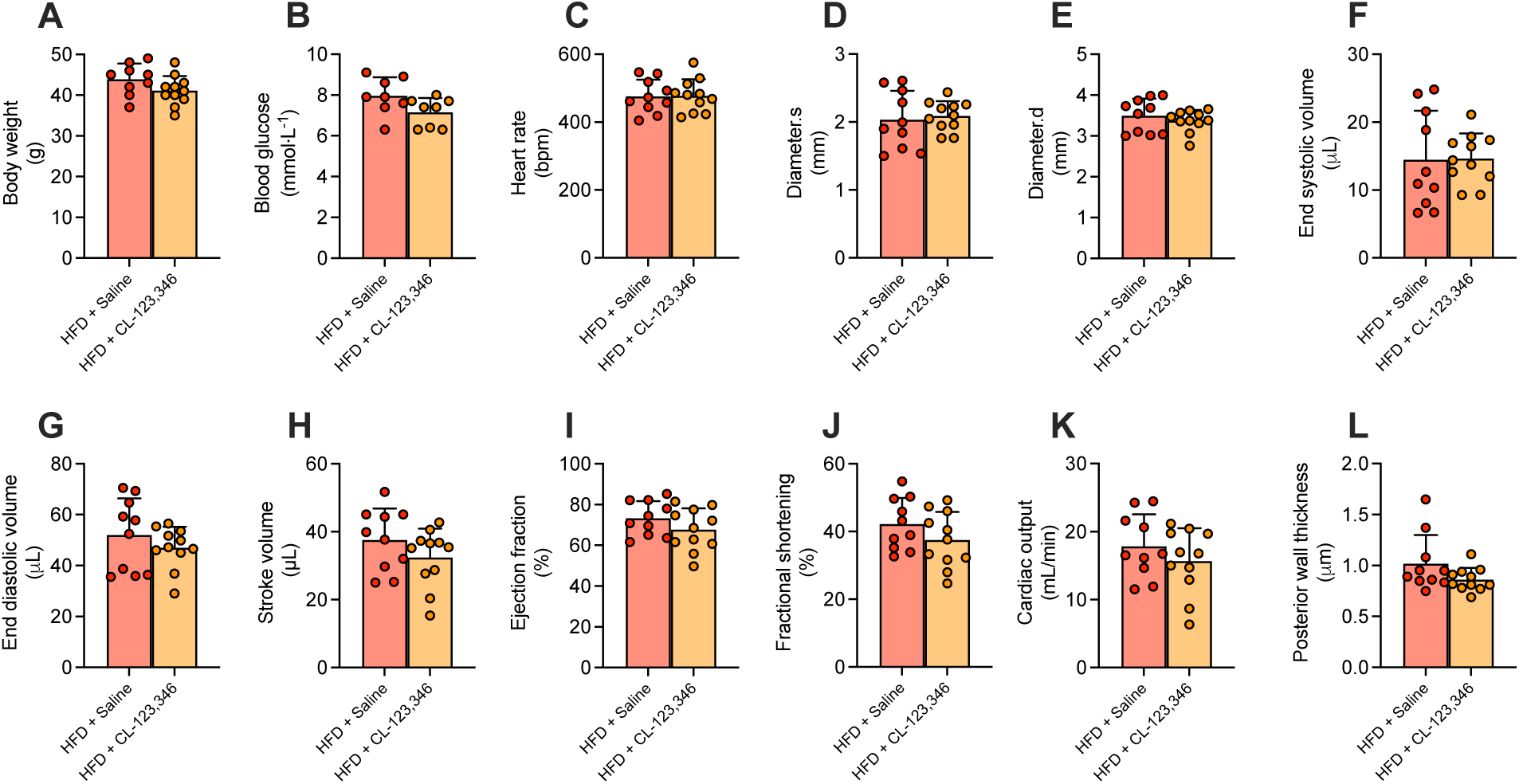
– Baseline comparison between HFD and HFD + CL- 123,346 groups. (A) body weight, (B) blood glucose, (C) heart rate, (D) LV diameter during systole and (E) diastole, (F) end systolic volume, (G) end diastolic volume, (H) stroke volume, (I) ejection fraction, (J) fractional shortening, (K) cardiac output, and (L) posterior wall thickness. Statistical analysis: two-tailed Student’s t-test. * p<0.05.

**Supplementary Figure 6.**
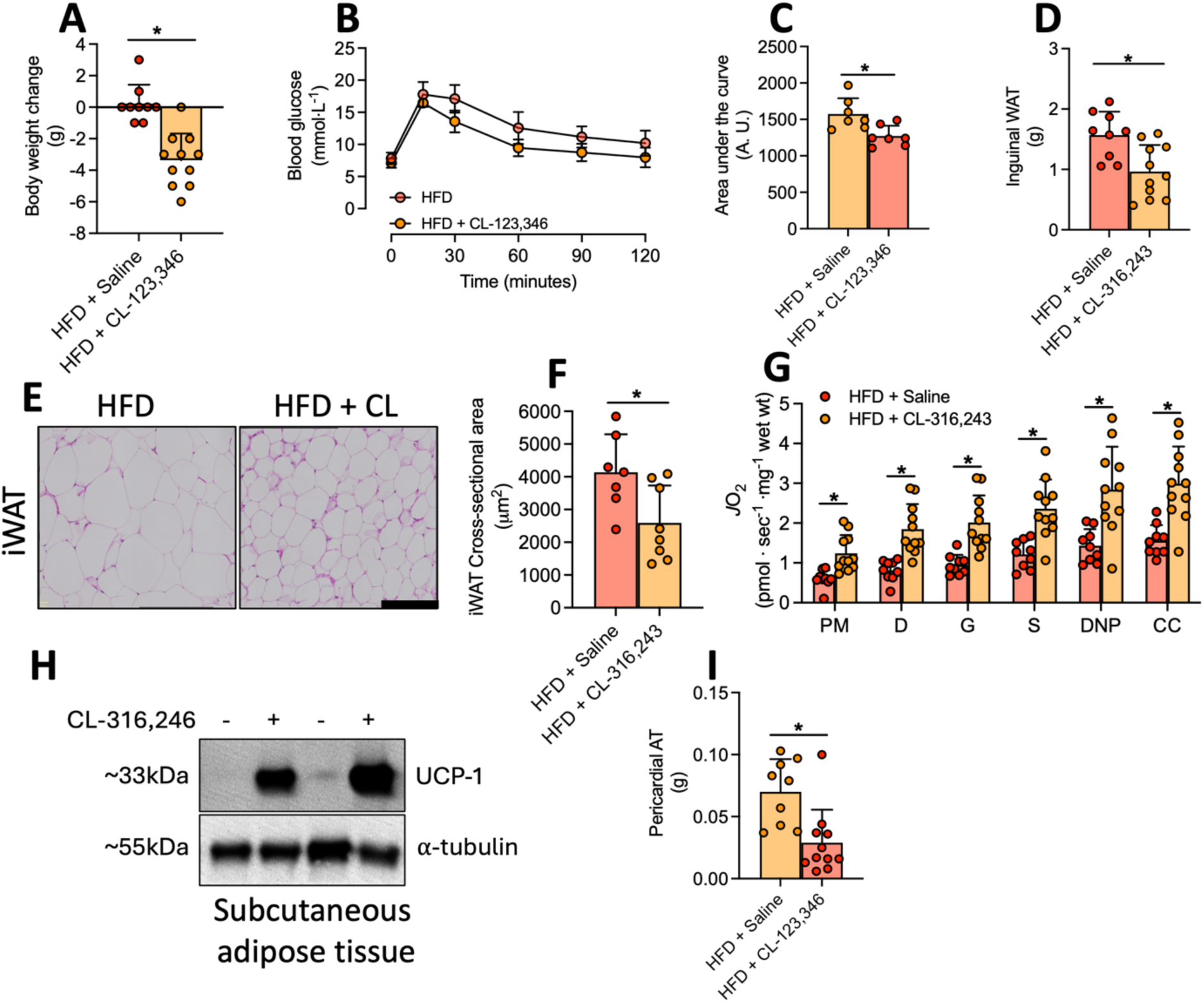
– Metabolic effects of CL-123,346 treatment in HFD-fed mice. (A) body weight change, (B) blood glucose excursion during ipGTT, and (C) area under the curve. (D) Inguinal WAT mass, (E) representative histological images of iWAT and (F) cross-sectional area. (G) Mitochondrial oxygen consumption and (H) representative western blot image of UCP1 protein. (I) pericardial AT mass. ipGTT – intraperitoneal glucose tolerance test; HFD – high-fat diet; WAT – white adipose tissue; P – pyruvate; M – malate; D – ADP; G – glutamate; S – succinate; DNP - 2,4-Dinitrophenol; CC – cytochrome c; UCP1 – uncoupling protein 1. Statistical analysis: two-tailed Student’s t-test. * p<0.05.

**Supplementary Figure 7.**
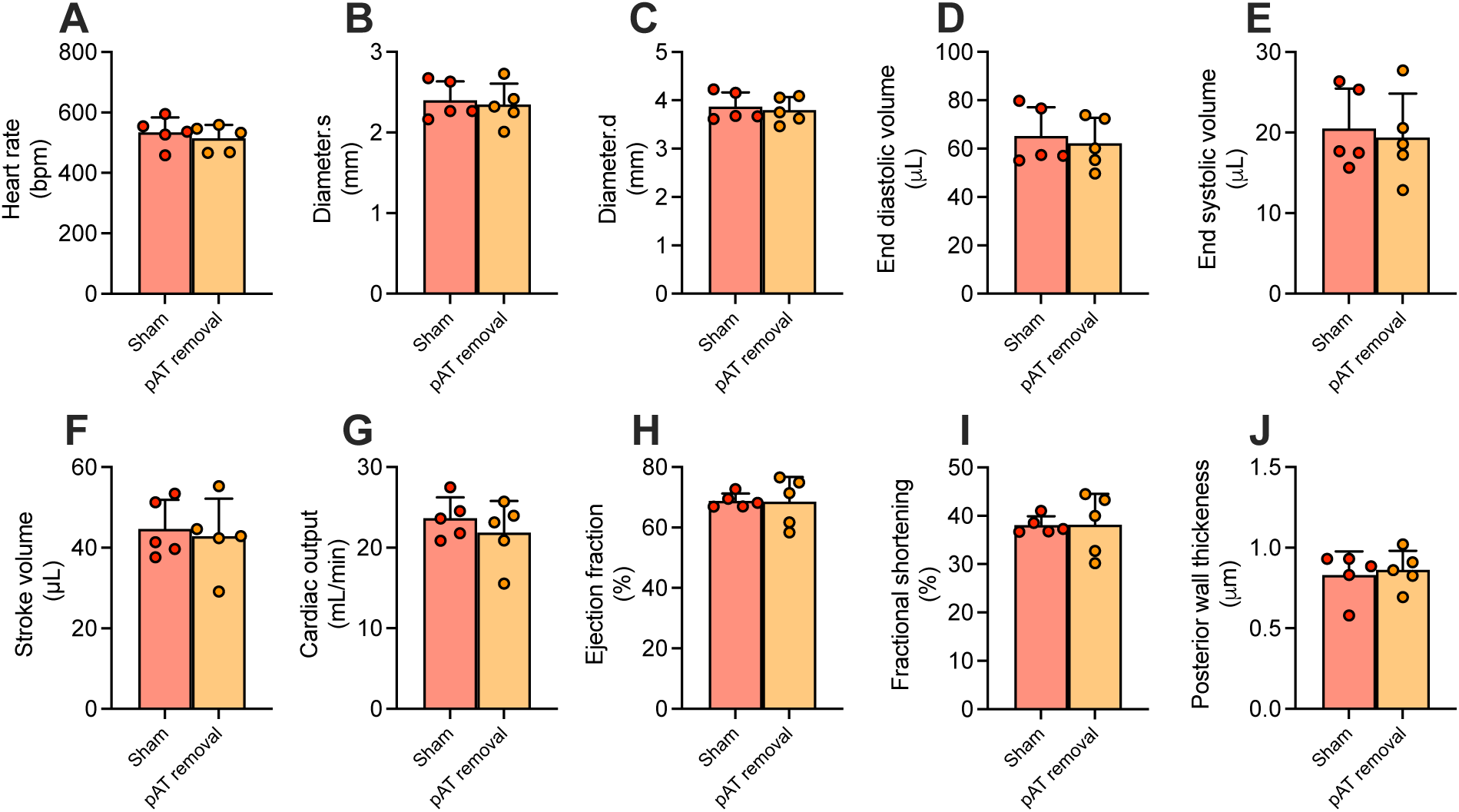
– Removal of pAT in lean mice does not affect heart function (A) Heart rate, (B) LV diameter during systole or (C) diastole, (D) end diastolic volume, (E) end systolic volume, (F) stroke volume, (G) cardiac output, (H) ejection fraction, (I) fractional shortening, and (J) posterior wall thickness during diastole. LV – left ventricle. Statistical test: two-tailed Student’s t-test.

**Supplementary Figure 8.**
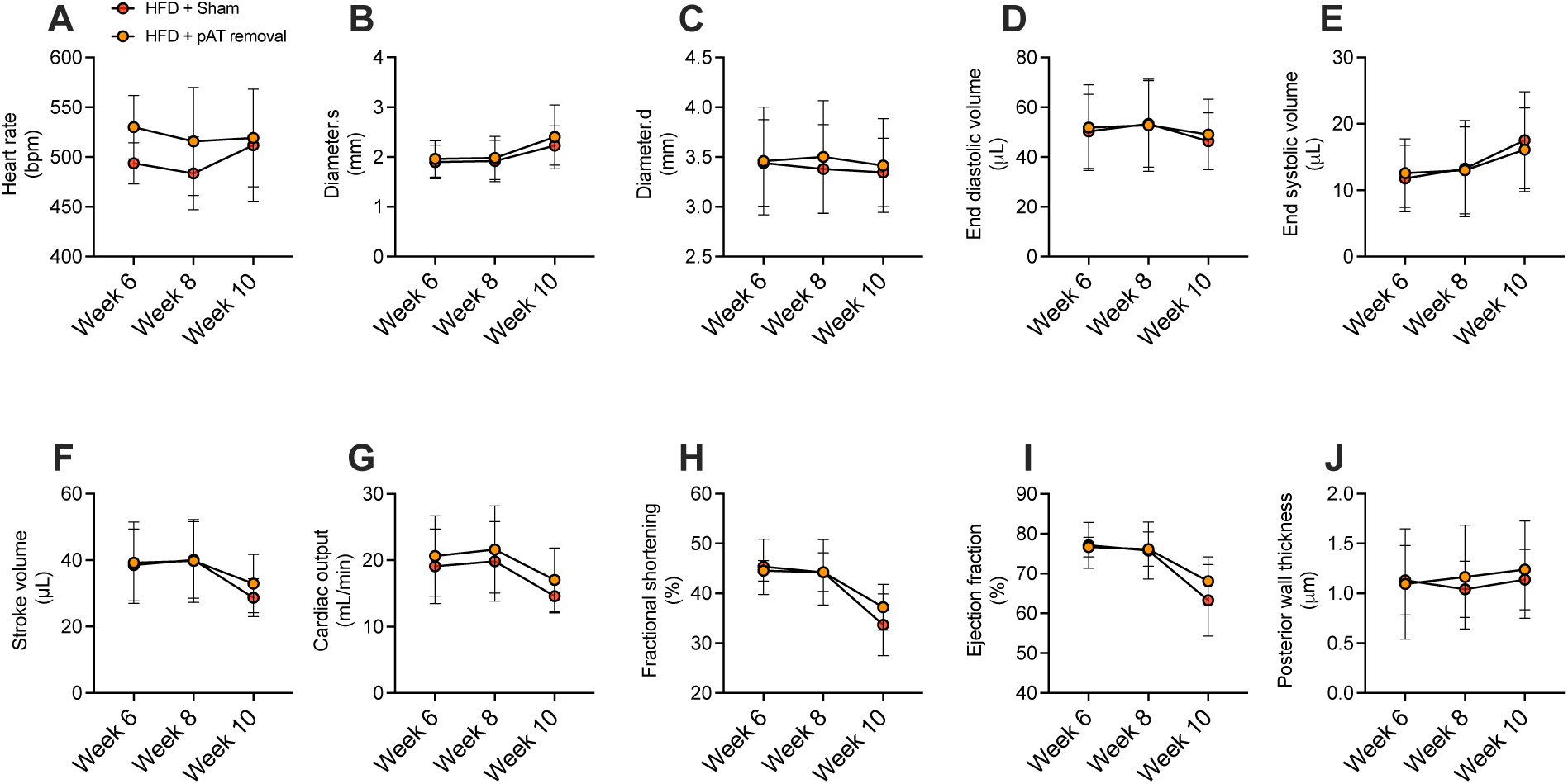
– Removal of pAT does not affect heart function during the development of obesity. (A) Heart rate, (B) LV diameter during systole or (C) diastole, (D) end diastolic volume, (E) end systolic volume, (F) stroke volume, (G) cardiac output, (H) fractional shortening, (I) ejection fraction, and (J) posterior wall thickness during diastole. LV – left ventricle. Statistical test: two-way ANOVA with Tukey’s post-hoc test.

**Supplementary Figure 9.**
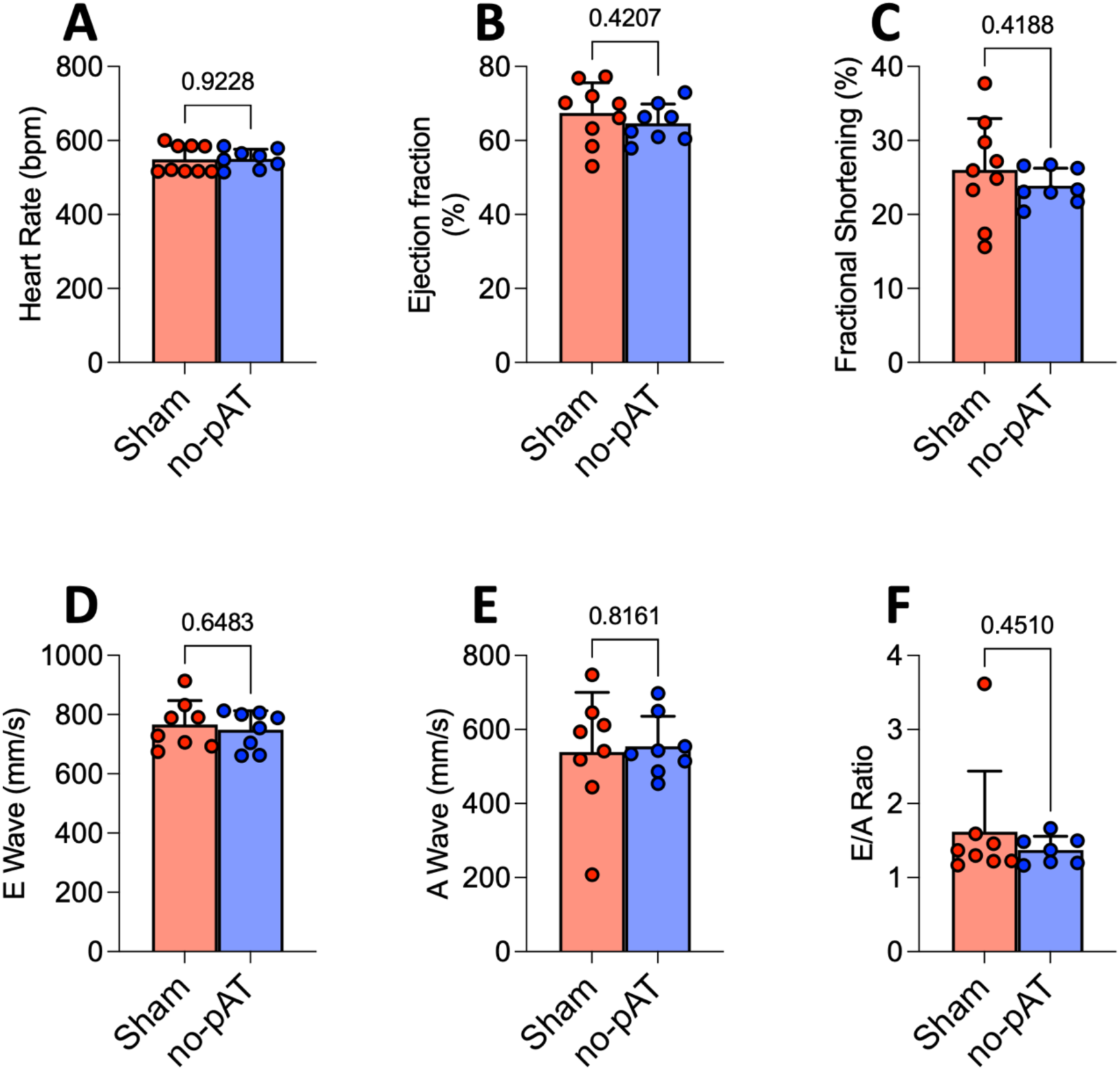
– Echocardiogram exams following pAT removal from obese mice. (A) Heart rate, (B) ejection fraction, (C) fractional shortening, (D) E wave, (E) A wave, and (F) E/A ratio. Sham – animals with no pAT removal. No-pAT – animals with pAT removed at 8^th^ week of HFD-feeding, evaluated 3 weeks later under the same diet. pAT – pericardial adipose tissue; HFD – high-fat diet. Statistical test: two-tailed Student’s t-test.

**Supplementary Figure 10.**
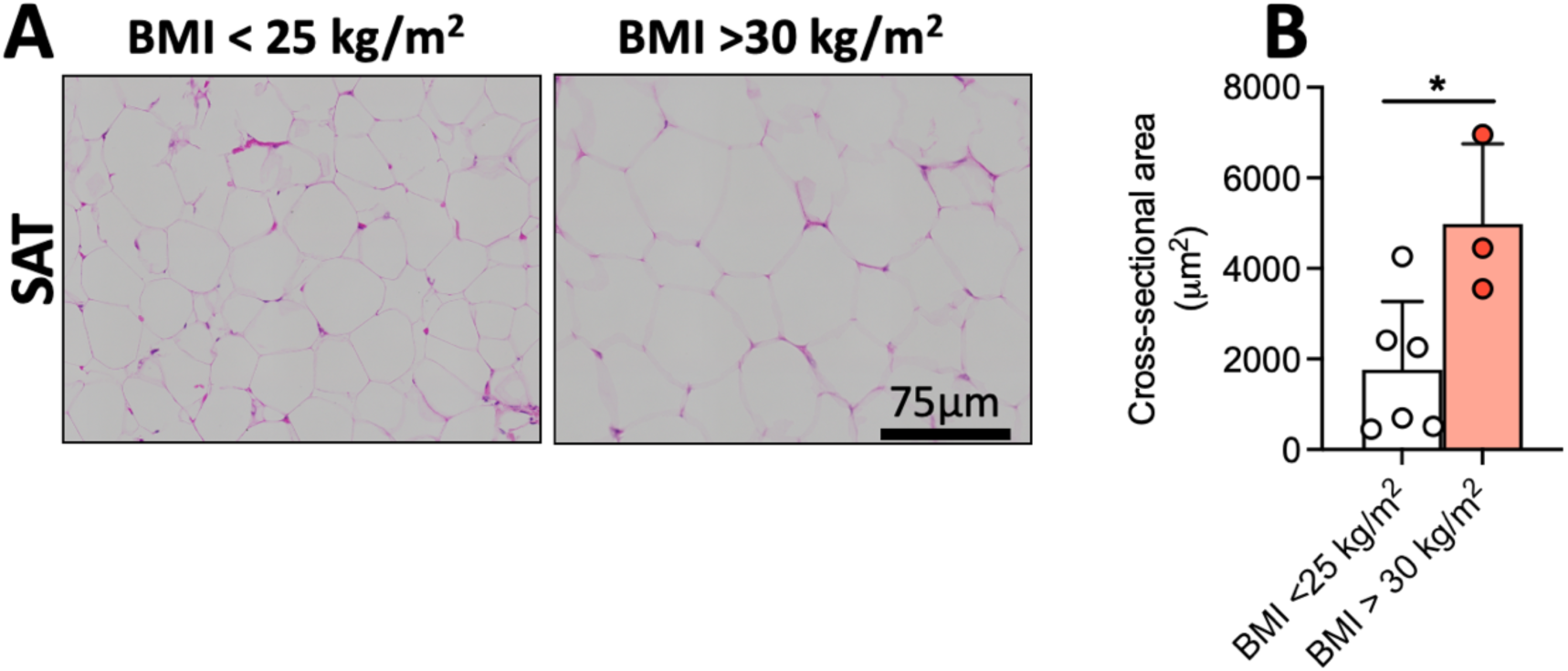
– Histology of subcutaneous adipose tissue from lean and obese humans. (A) Representative images of white subcutaneous adipose tissue from lean and obese and (B) quantification of average cross-sectional area of adipocytes. SAT – subcutaneous adipose tissue; BMI – body mass index. Statistical analysis: two-tailed Student’s t-test. * p<0.05.

